# Cryo-EM structures of inactive and G_i_-coupled GABA_B_ heterodimer

**DOI:** 10.1101/2020.04.21.053488

**Authors:** Chunyou Mao, Cangsong Shen, Chuntao Li, Dan-Dan Shen, Chanjuan Xu, Shenglan Zhang, Rui Zhou, Qingya Shen, Li-Nan Chen, Zhinong Jiang, Jianfeng Liu, Yan Zhang

## Abstract

Metabotropic GABA_B_ G protein-coupled receptor functions as a mandatory heterodimer of GB1 and GB2 subunits and mediates inhibitory neurotransmission in the central nervous system. Each subunit is composed of the extracellular Venus flytrap (VFT) domain and transmembrane (TM) domain. Here we present cryo-EM structures of human full-length heterodimeric GABA_B_ receptor in the antagonist-bound inactive state and in the active state complexed with agonist and positive allosteric modulator in the presence of G_i1_ protein at a resolution range of 2.8-3.0 Å. Cryo-EM analysis of the activated-GABA_B_–G_i1_ complex revealed that G_i1_ couples to the activated receptor primarily in three major conformations, one via GB1 TM and two via GB2 TM, respectively. Our structures reveal that agonist binding stabilizes the closure of GB1 VFT, which in turn triggers a rearrangement of TM interfaces between two subunits from TM3-TM5/TM3-TM5 in the inactive state to TM6/TM6 in the active state and finally induces the opening of intracellular loop 3 and synergistically shifting of TM3, 4 and 5 helices in GB2 TM domain to accommodate the α5-helix of G_i1_. These results provide a structural framework for understanding class C GPCR activation and a rational template for allosteric modulator design targeting dimeric interface of GABA_B_ receptor.

## Introduction

Metabotropic GABA_B_ receptor is a G protein-coupled receptor (GPCR) for the major inhibitory neurotransmitter, γ-amino butyric acid (GABA), in the central nervous system to mediate slow and prolonged inhibitory activity^1-3^. GABA_B_ receptor couples to G_i/o_ proteins to inhibit neurotransmitter release in pre-synaptic neurons and cause hyperpolarization in the post-synaptic neurons^4^. GABA_B_ receptor suppresses adenylyl cyclase through Gα_i/o_, while gating ion channels and transactivating receptor tyrosine kinases via Gβγ^5-8^. Dysfunctions of GABA_B_ receptor or mutation in its genes have been implicated in a variety of neurological and psychiatric disorders including epilepsy, pain, anxiety, depression, schizophrenia, drug addiction, Rett syndrome and epileptic encephalopathies^4,9^. Recent studies reveal that the auto-antibodies of GABA_B_ receptor are possibly the origin of epilepsies and encephalitis^10,11^. Although a large number of antagonists and agonists and positive or negative allosteric modulators (PAMs or NAMs) have been developed for GABA_B_ receptor^12^, only two agonists approved as therapeutic drugs: baclofen (Lioresal®) used for the treatment of muscle spasticity and alcohol addiction^13,14^ and γ-hydroxybutyrate (GHB) used for the treatment of narcolepsy^15^.

GABA_B_ receptor belongs to the class C GPCR composed by metabotropic glutamate receptors (mGlus), calcium-sensing receptor (CaSR) and taste 1 receptors^16^. Class C GPCR form obligatory dimers, among them, mGlus and CaSR form the homodimer^17,18^, whereas GABA_B_ receptor is an obligatory heterodimer of two subunits GABA_B1_ (GB1) and GABA_B2_ (GB2)^19,20^. Each subunit is composed of a large extracellular ‘Venus Flytrap’ (VFT) domain, a transmembrane (TM) domain, and a cytoplasmic tail^21^ (Extended Data Fig. 1a, b). GB1 possesses an endoplasmic reticulum (ER) retention motif in its cytoplasmic tail^22^. The interaction between GB1 and GB2 facilitates its cell surface expression through coiled-coil interactions in its cytoplasmic tail^23^. While GB1 is responsible for ligand recognition through its VFT^24^, GB2 couples G_i/o_ proteins through its TM^25,26^. However, a line of evidence show that in the absence of GB2, GB1_asa_ (a GB1 mutant with deletion of ER retention signal) at the cell surface or ER-localized GB1 also induces downstream signaling through G_i/o_ proteins^27,28^.

**Extended Data Fig.1.**
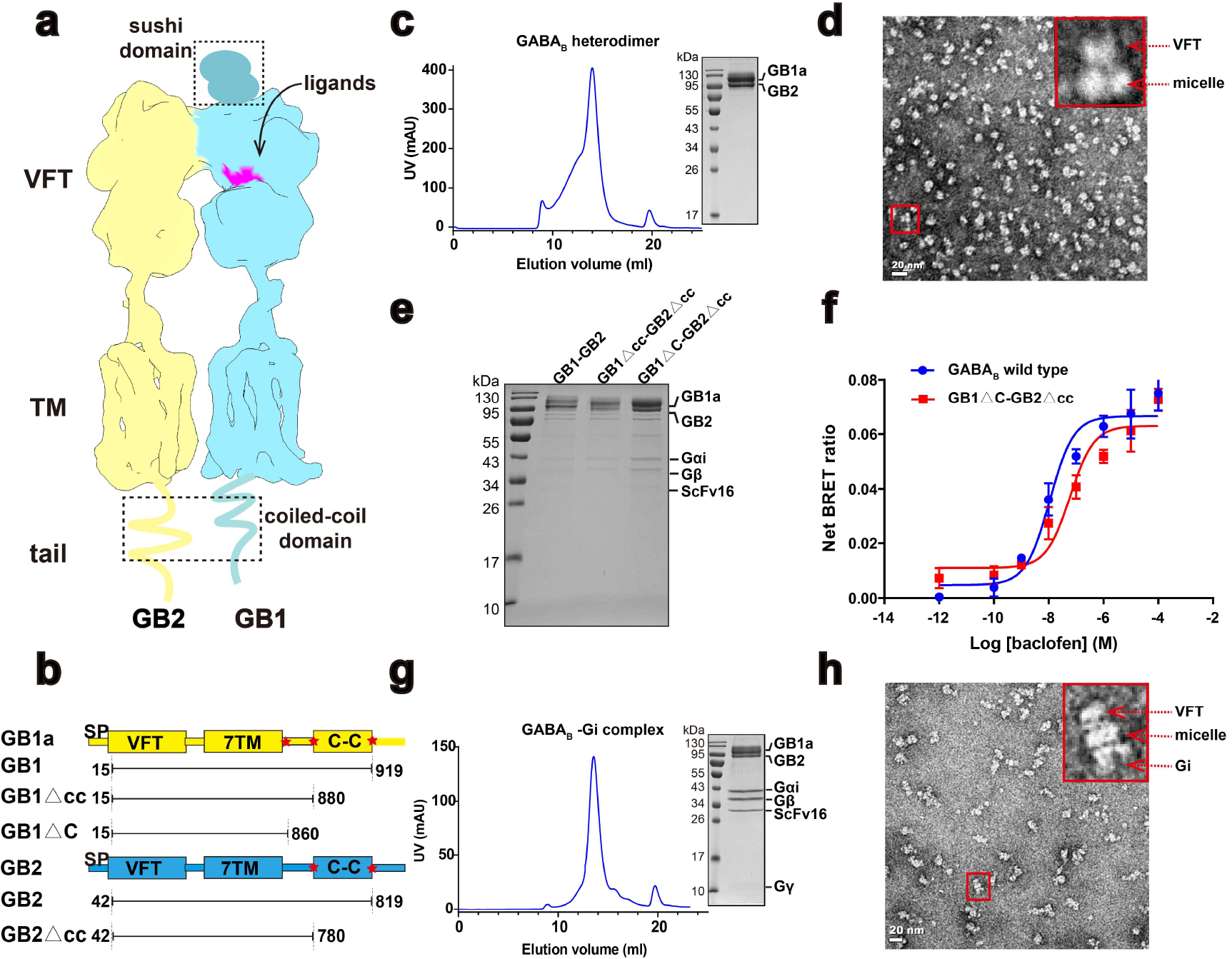
Purification and characterization of the GABA_B_ heterodimer and GABA_B_–G_i1_ complex. **a**, Schematic of the heterodimeric GABA_B_ receptor. **b**, Schematic of constructs used in this study. △cc represents the coiled-coil (c-c) domain is removed. △C represents the entire C-terminus after TM domain in GB1 or GB2 is truncated. **c**, Size-exclusion chromatography profile and SDS-PAGE gel of the purified antagonist-bound GABA_B_ receptor. **d**, Negative staining EM analysis of the purified inactive GABA_B_ receptor shows the clear density for VFT and detergent micelles. **e**, G protein pull-down analysis of the different truncations (GB1-GB2, GB1△cc-GB2△cc, and GB1△C-GB2△cc) of GABA_B_ receptor. SDS-PAGE gel showed that GB1△C-GB2△cc exhibits a greater efficiency in complex formation among these constructs and were selected for further structural study. **f**, Dose-response curve of baclofen-induced BRET changes in the Gαi sensors for the wide-type and GB1△C-GB2△cc construct of GABA_B_ receptor, showing that GB1△C-GB2△cc exhibits similar phrenology to that of wild-type receptor. **g**, Size-exclusion chromatography profile and SDS-PAGE gel of the purified agonist/PAM bound GABA_B_–G_i1_ complex using GB1△C-GB2△cc construct. **h**, Negative staining EM visualization of the purified GABA_B_-G_i1_ complex showing clear density for VFT, detergent micelles and G_i1_ protein.

The crystal structures of isolated heterodimeric VFT of GABA_B_ receptor in presence of the antagonist or agonist revealed the open or closed conformation of GB1 VFT^29^. Structures of isolated VFT and TM domains from other class C GPCR have been reported^30-32^, in addition to more recently breakthrough that cryo-EM structures of full-length mGlu5 in apo state and in the presence of agonist have been determined both at overall 4 Å resolution, providing the first insights into the architecture of mGlu5 and structural framework of mGlu5 activation^33^. However, no structure of full-length GABA_B_ receptor has been solved, limiting our understanding of the configuration of GABA_B_ receptor. Moreover, molecular mechanism of class C GPCR signal transduction remains elusive, primarily owning to the lack of structural information of the receptor in various states, especially coupled to a down-streaming G protein. Here we present cryo-EM structures of human full-length heterodimeric GABA_B_ receptor in the antagonist-bound inactive state and in the agonist/PAM-bound active state complexed with heterotrimeric G_i1_ protein, shedding lights on class C GPCR activation and providing a rational template for allosteric modulator design targeting dimeric interface of GABA_B_ receptor.

## Results

### Cryo-EM structure determination of GABA_B_ receptor

We first sought to obtain the heterodimeric human GABA_B_ receptor in the fully inactive state, thus we overexpressed both GB1a and GB2 subunits in full length in mammalian cells and maintained 20 µM CGP54626, a potent antagonist with estimated nano-molar affinity to the receptor^34^, through the sample preparation stage. The heterodimeric GABA_B_ receptor was solubilized from the membrane using detergent. The following visualization by negative-staining EM displayed the existence of the intact heterodimer, and unlike detergent-reconstituted mGlu5 in the inactive (apo) state with split micelles^33^, GABA_B_ receptor showed one large disc-shaped micelle containing TMs from both subunits (Extended Data Fig. 1c, d). Two-dimensional class averages of frozen specimen showed clear secondary structure features for GABA_B_ receptor dimer (Extended Data Fig. 2), confirming two TM bundles embedded in the detergent micelle suggesting the more stable intersubunit interactions between TM domains of GABA_B_ receptor compared to that of mGlu5^33^. We therefore obtained cryo-EM images and determined the density map for CGP54626-bound GABA_B_ receptor using single particle cryo-EM to an overall resolution of 3.0 Å. Local resolution calculations indicate a range of 2.4-3.4 Å in most map regions (Fig. 1a, Extended Data Fig. 2). The antagonist, VFT and TM domains from both subunits are clearly visible in the cryo-EM map (Fig. 1a), except the sushi domains at GB1 N terminus and coiled-coil domain at the GABA_B_ receptor cytoplasmic tail, indicating the dynamic nature of these two regions.

**Fig.1.**
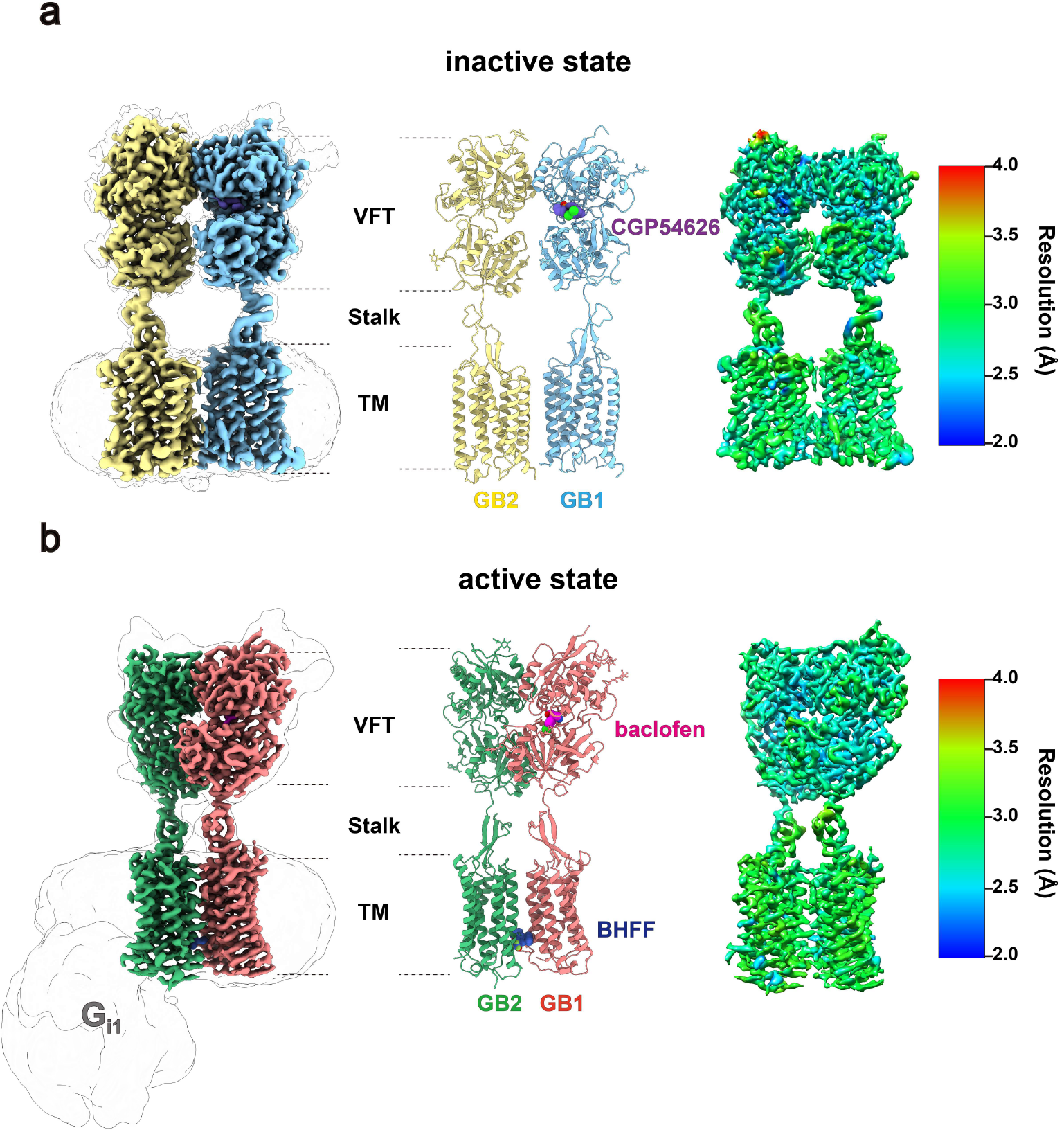
Cryo-EM structures of GABA_B_ receptor in the inactive and active states. **a-b**, Cryo-EM density maps (left), models (middle) and maps coloured according to local resolution (right) for the GABA_B_ receptor in the presence of CGP54626 (slate; antagonist) (**a**) and in complex with baclofen (magenta; agonist), BHFF (steel blue; PAM) and G_i1_ protein (**b**). The coloured density map is a composite map generated with the VFT and TMD locally refined map. The cryo-EM density map before focused refinement in transparent superposed with the final map, illustrating density for detergent micelle and G_i1_ protein. The local resolution (Å) was calculated with the locally refined map as input, indicating a range of 2.5-3.5 Å resolution in most map regions for both the inactive and active states. GB1 and GB2 in inactive state, blue and yellow, respectively; red and green in active state, respectively.

**Extended Data Fig.2.**
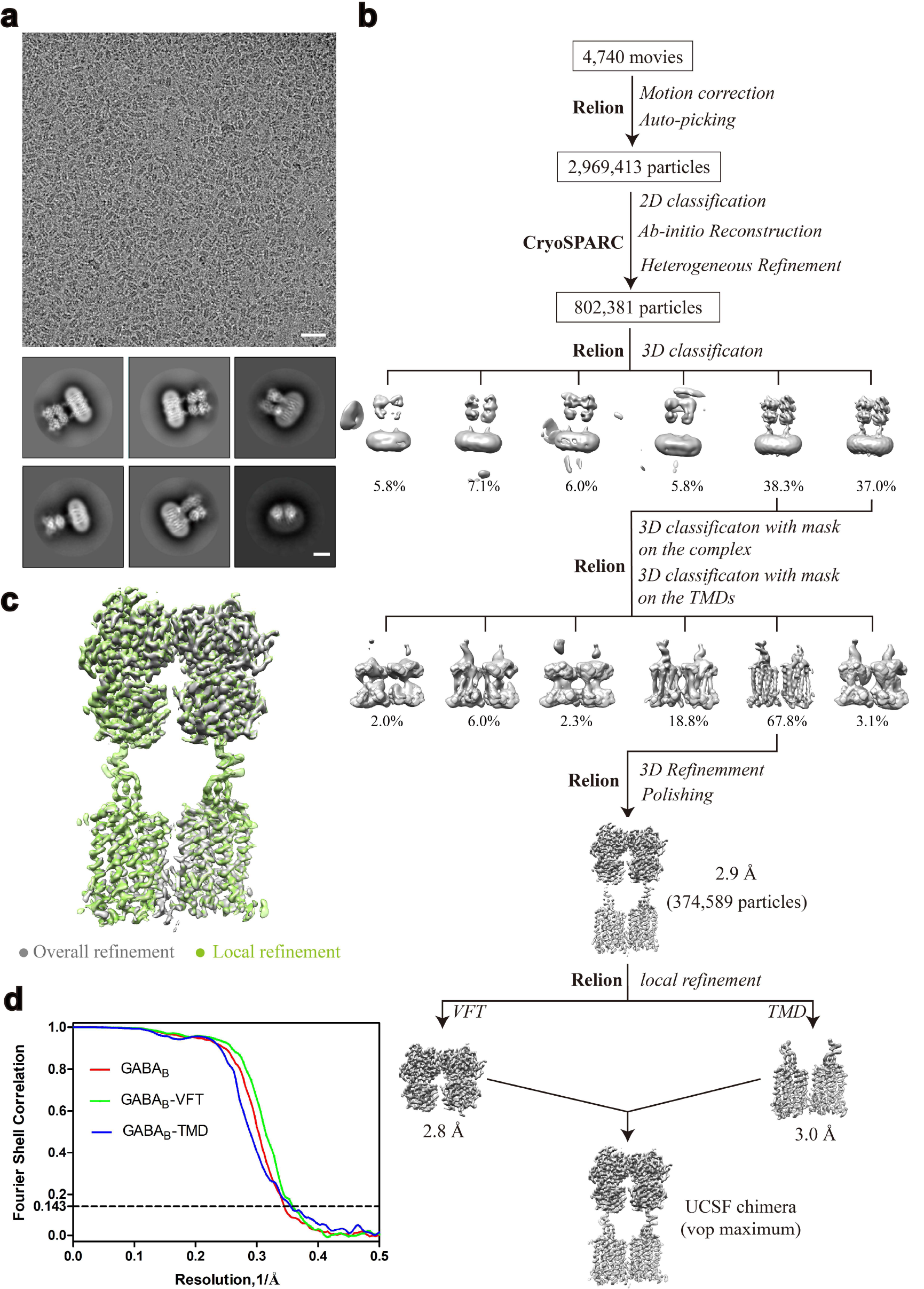
Single-particle cryo-EM of the CGP54626-bound GABA_B_ receptor. **a**, Cryo-EM micrograph (scale bar: 30 nm) and representative two-dimensional class averages of the inactive GABA_B_ receptor in LMNG detergent micelles (scale bar: 5 nm). **b**, Flow chart of cryo-EM data processing. Details are described in Method section. **c**, Comparison of the overall refined map and the locally refined composite map. The locally refined composite map exhibits a slight improvement of the TMs density and adopts almost the identical conformation with the overall refined map. **d**, Gold-standard Fourier shell correlation (FSC) curves of overall refined receptor and the locally refined VFT and TMD.

To prepare the stable and sufficient activated GABA_B_–G_i1_ complex for cryo-EM study, initial efforts were made to accomplish the complex formation under the condition in which the receptor and the purified G_i1_ protein were incubated with baclofen, a commercially available PAM BHFF in racemate form (rac-BHFF)^35^ and G_i_-binding protein scFv16^36^, unfortunately leading to the oligomerization of the receptor (data not shown). Considering the flexibility of the coiled-coil domain indicated by the inactive structure of the full-length receptor, we thus removed it and pull-down assay suggested that the receptor lacking of the coiled-coil domain (GABA_B_Δcc) exhibited increased expression level and improved complex formation yield of GABA_B_–G_i1_ (Extended Data Fig. 1e). These modifications do not alter receptor pharmacology (hereafter referring GABA_B_Δcc as GABA_B_ unless noted otherwise) (Extended Data Fig. 1f). We therefore formed the complex between the GABA_B_ in mammalian cells, heterotrimeric G_i1_ protein in insect cells before solubilization of the baclofen/PAM–GABA_B_–G_i1_ complex using lauryl maltoseneopentyl glycol and purification by tandem affinity and size-exclusion chromatography (Extended Data Fig. 1g). Sample evaluation by negative-stain EM confirmed the presence of G_i1_ protein and monodispersed particle distribution (Extended Data Fig. 1h). In addition, cryo-EM analysis suggested multiply conformations spontaneously present in the GABA_B_–G_i1_ complex and three-dimensional classification revealed three major conformers, with one of which representing G_i1_ protein engaged with GB1 and two of which representing G_i1_ protein coupled to GB2 in distinct orientations (termed B1, B2a and B2b state, respectively). We further refined these three conformers independently and obtained the reconstructions with nominal resolutions of 8.8 Å, 6.8 Å and 8.6 Å, respectively, which were adequate for rigid-body docking of the receptor and Gi1 independently to generate models for their relative arrangement (Extended Data Fig. 3). Close visualization and superimposition of three conformers with each other revealed no distinguishable difference in the receptor part, enabling us to obtain high-resolution reconstruction for the active receptor (Extended Data Fig. 3, 4). Therefore, we focused particle projection alignment on the VFT module and TM domain of the receptor separately to improve the map quality and managed to obtain the maps with the resolutions of 2.9 Å and 3.3 Å, respectively, which were subsequently combined to produce a composite map for model building and refinement (Fig. 1b, Extended Data Fig. 3).

**Extended Data Fig.3.**
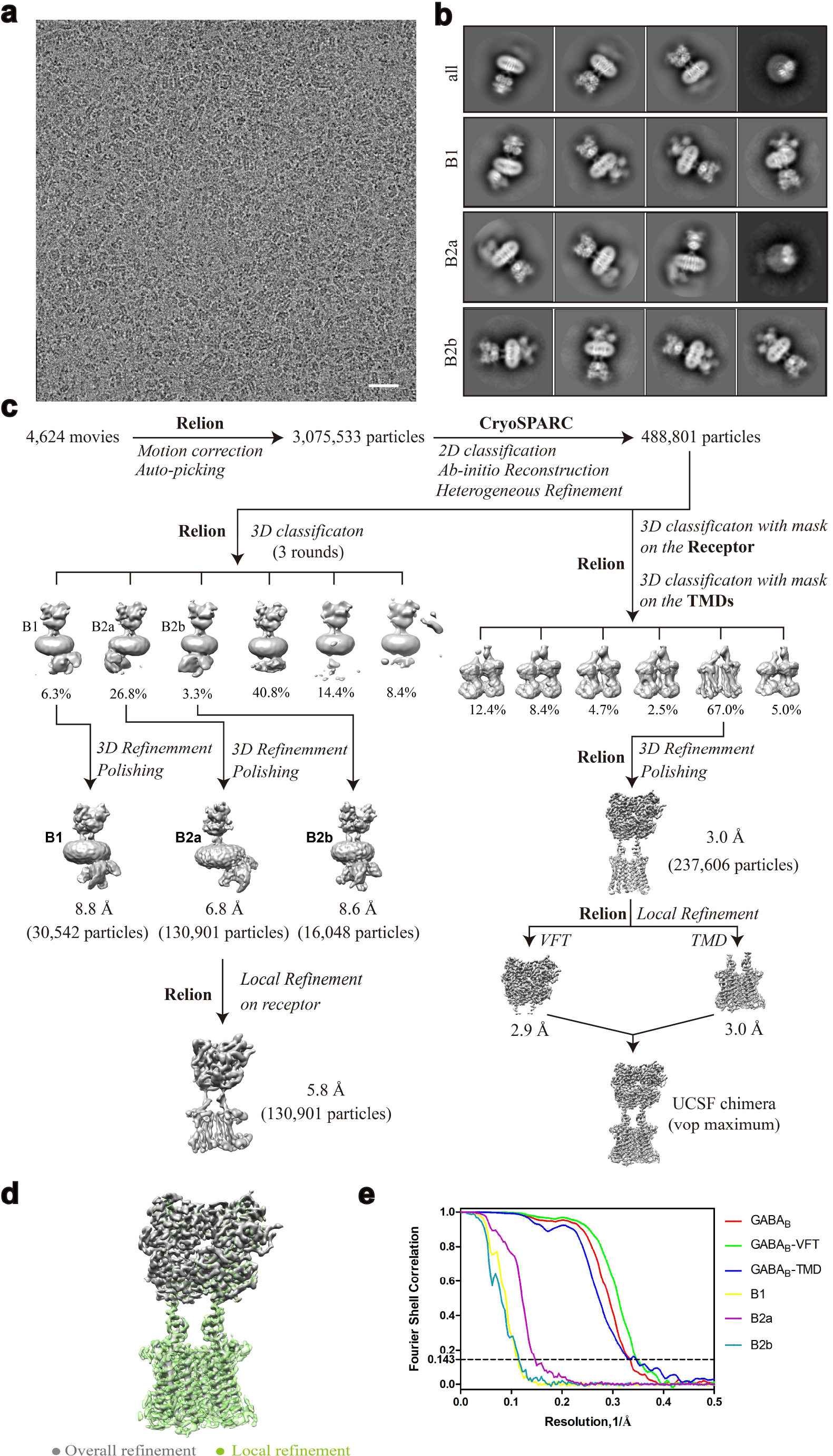
Single-particle reconstruction of the agonist/PAM-bound GABA_B_–G_i1_ complex. **a**, Representative cryo-EM micrograph of the GABA_B_–G_i1_ complex (scale bar: 30 nm). **b**, Representative two-dimensional class averages of the overall refined GABA_B_ receptor (all) and the GABA_B_–G_i1_ complex in B1, B2a and B2b states, respectively. (scale bar: 5 nm). **c**, Flow chart of cryo-EM data processing. Details are described in method section. **d**, Comparison of the overall refined map and the locally refined composite map. The locally refined composite map shows a substantial improvement for the TM density and exhibits almost the same conformation with the overall refined map. **e**, Gold-standard Fourier shell correlation curves of overall refined receptor, the locally refined VFT and TMD and the refined GABA_B_-G_i1_ complex in B1, B2a, B2b states.

**Extended Data Fig.4.**
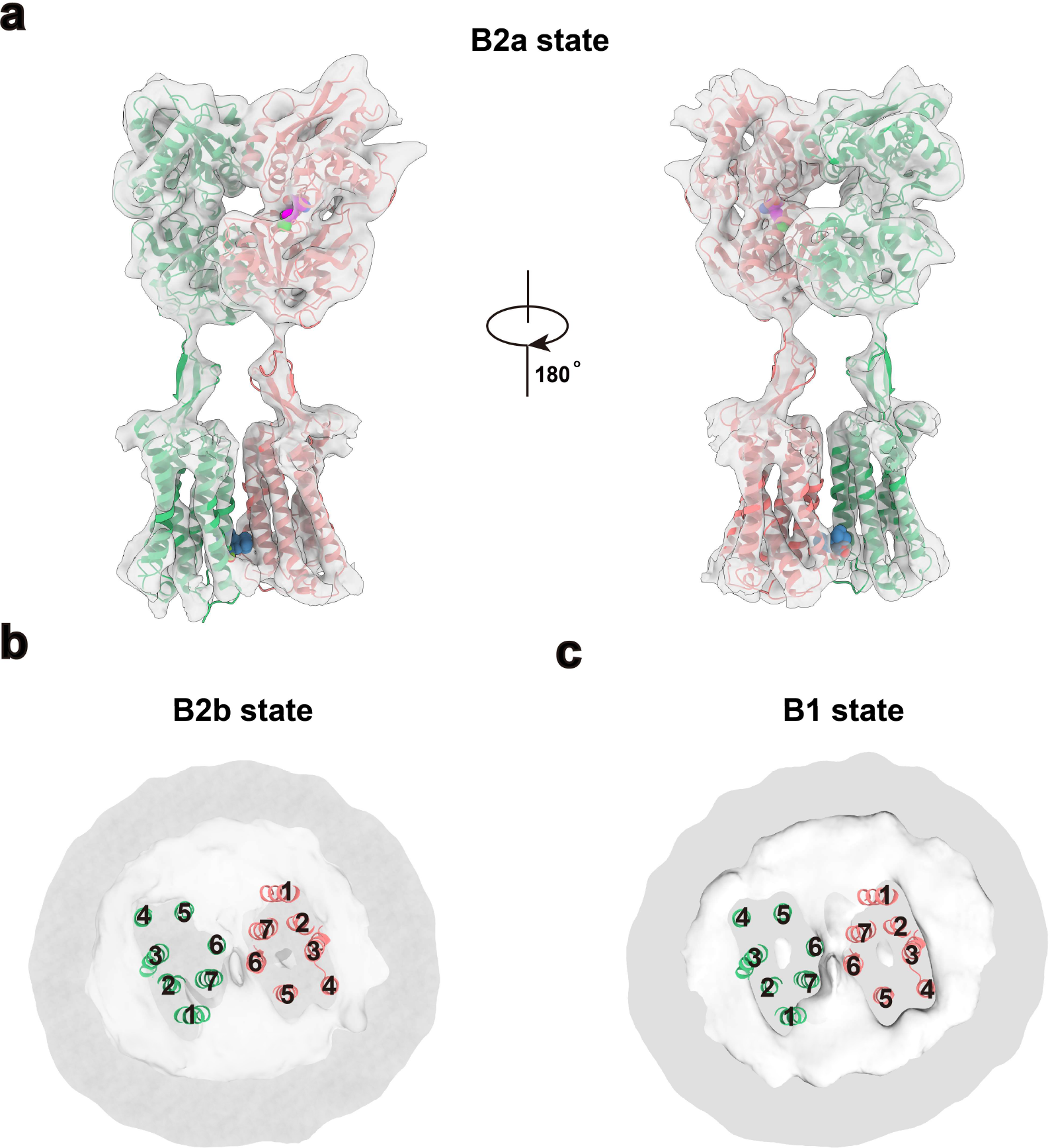
Fitting of the active GABA_B_ structure into the cryo-EM density maps of GABA_B_–G_i1_ complex in B2a, B2b and B1 states. **a**, Side views of the active GABA_B_ structure derived from the high-resolution density map against cryo-EM map of GABA_B_–G_i1_ complex in B2a state. **b-c**, Cut through of bottom-up views of the active GABA_B_ structure derived from the high-resolution density map against cryo-EM map of GABA_B_–G_i1_ complex in B2b state (**b**) and B1 state (**c**), respectively. The model of GABA_B_ receptor was rigid-body docked into the density.

Taken together, the high-quality density maps allow the unambiguously placement of antagonist (CGP54626), agonist (baclofen), PAM (BHFF) and most side chains of receptor amino acids in both inactive and active states including all the extracellular loops (ECLs) and intracellular loops (ICLs) with the exception of ICL2 (Fig. 3e, Extended Data Fig. 5, 7). Thus, our structures provide accurate models of the full-length GABA_B_ receptor in the inactive and active states, shed lights on intersubunit interactions between GB1 and GB2, ligand recognition and proposed models of G_i1_ engagement to the receptor. We also observed multiply ordered cholesterols not only surrounding the periphery of both TM domains like many other GPCRs^37,38^, but also between two subunits primarily in the inactive state (Extended Data Fig. 6a). Surprisingly, well-defined density with the shape of phospholipid in the TM bundle of each subunit was also observed in both states (Extended Data Fig. 6b-e). Considering the extensive interactions between each phospholipid and the corresponding subunit and the consistent presence of the phospholipids in both states, we speculated that phospholipids in the core pockets are more likely structural components rather than allosteric modulators.

**Fig.2.**
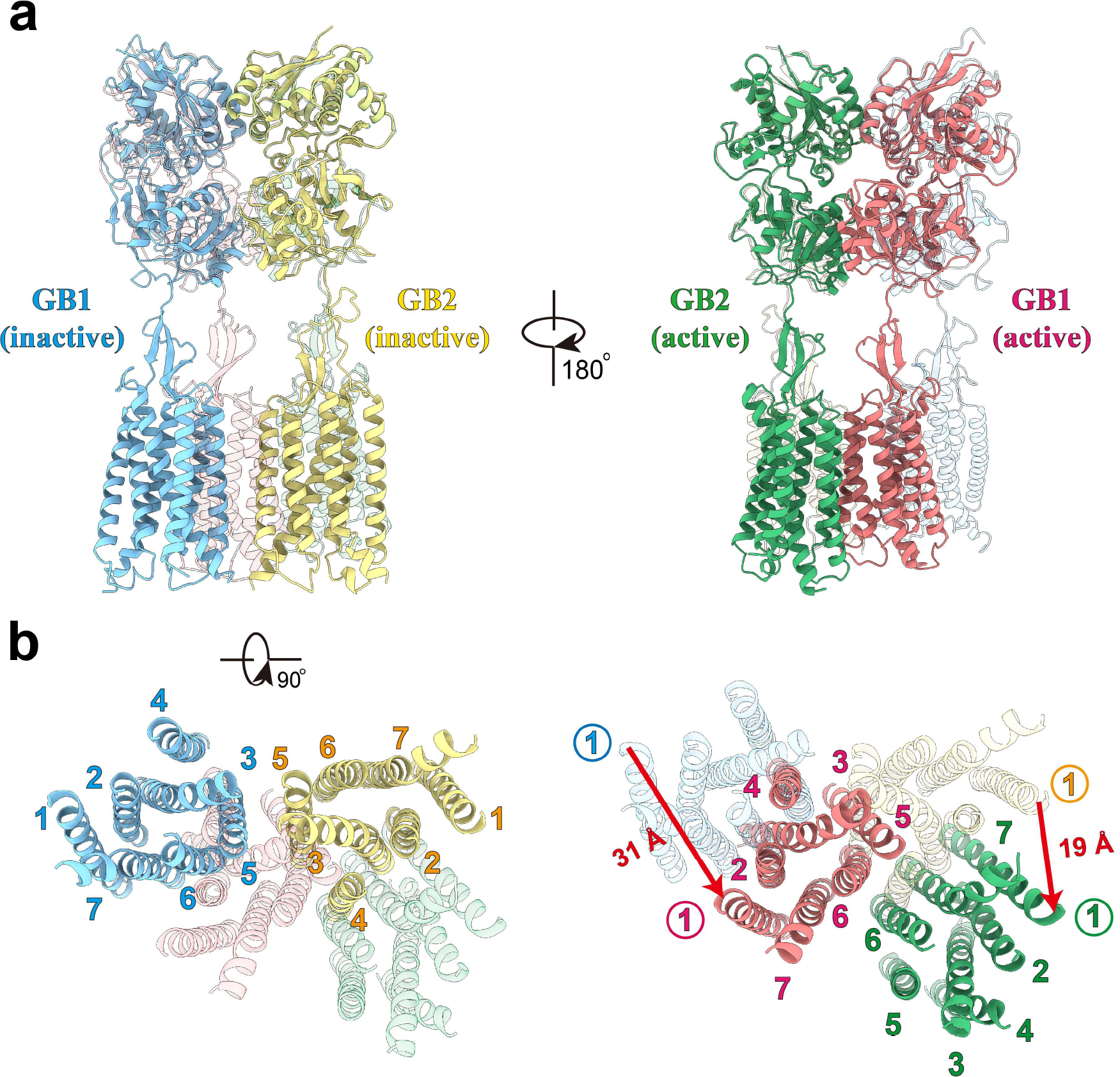
Structural comparison of GABAB receptor in inactive and active states. **a-b**, Orthogonal views of the superimposed structures of GABA_B_ receptor in inactive and active states, showing the domain repositioning upon agonist binding induced activation. Side views (**a**) and intracellular views (**b**) of superposed structures are shown, with the active structure in translucent in the left panels and the inactive structure in translucent in the right panels, respectively. VFT domains and loops are omitted for clarity in **b**. Red arrows indicate the translation direction and distance for GB1 and GB2 (measured at extracellular tips of TM1 helices), respectively. Structures were aligned on the combined domains of GB1 VFT and GB2 lobe1, the relatively stable parts of the receptor along activation pathway.

**Fig.3.**
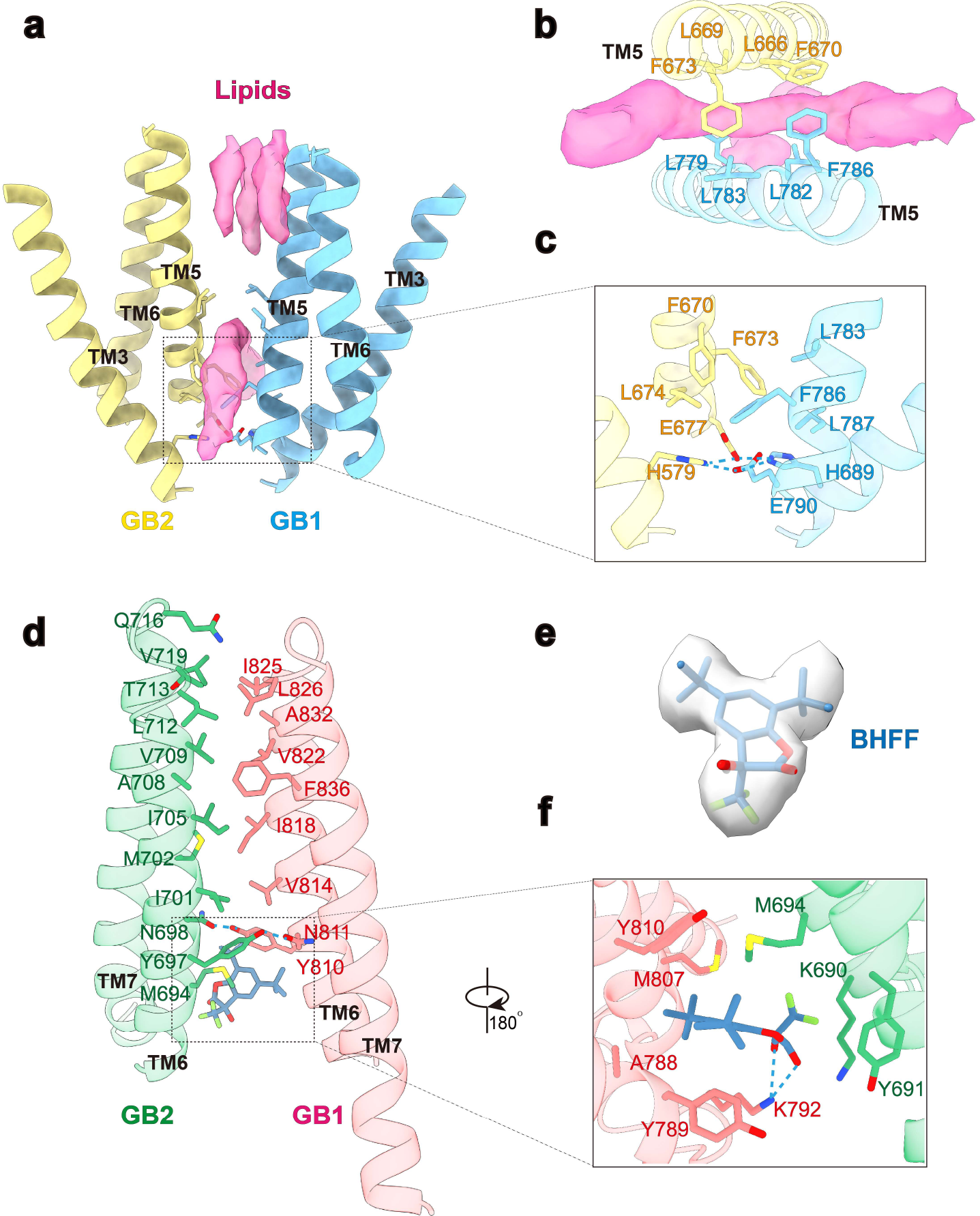
Structural details of the TM heterodimeric interfaces for the inactive and active GABA_B_ structures. **a-b**, Side (**a**) and intracellular (**b**) views of the inactive TM interface. EM densities (magenta) corresponding to putative cholesterols are shown in surface. **c**, close view of inactive TM interface in the proximal to the cytoplasm. H689^3.55^ and E790^5.60^ of GB1, H579^3.55^ and E677^5.60^ of GB2 forms salt bridge network to stabilize an inactive conformation. **d**, Detailed interactions of the TM interface in active state. **e-f**, Cryo-EM density (**e**) and the detailed interactions (**f**) of (+)-BHFF (PAM) in the active TM interface. The critical residues are shown as sticks. Hydrogen bonds are depicted as dashed lines.

**Extended Data Fig.5.**
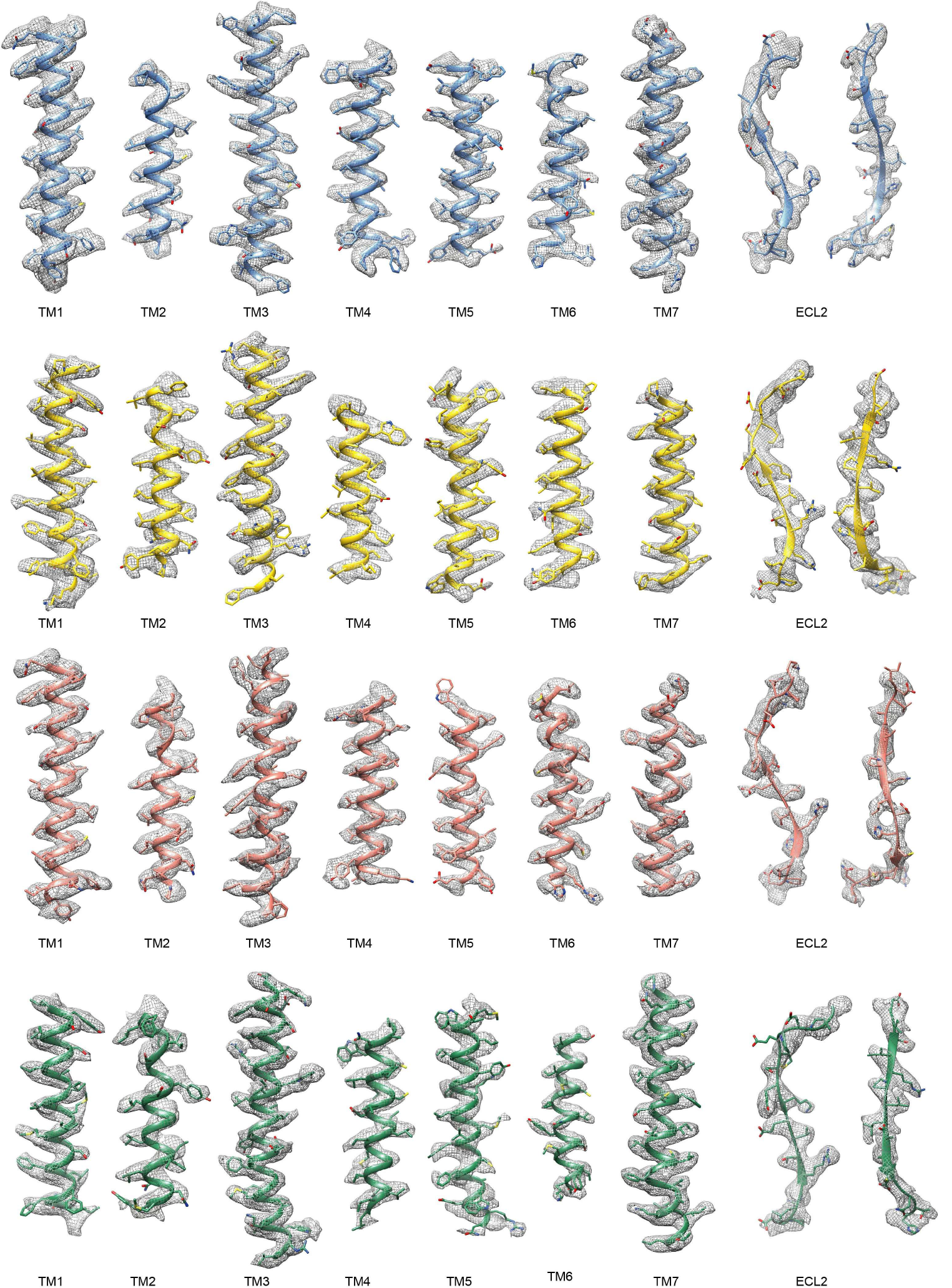
Electron density of transmembrane helices and ECL2 in GABA_B_ receptor. Cryo-EM density of transmembrane helices of GABA_B_ in inactive and active structures. Inactive GB1 in blue; Inactive GB2 in yellow; Active GB1 in red; Active GB2 in green.

**Extended Data Fig.6.**
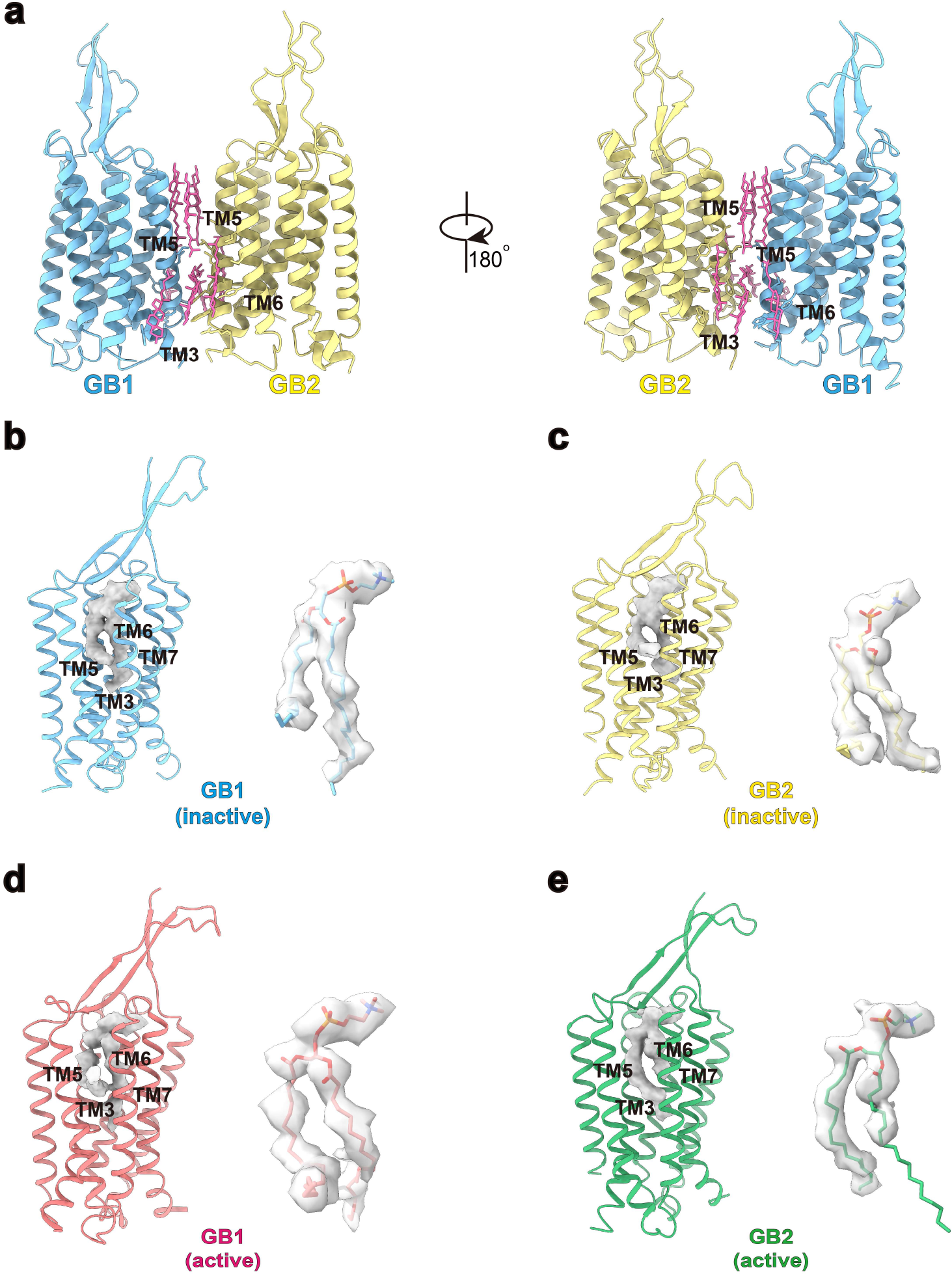
Putative cholesterols and phospholipids in GABA_B_ receptor. **a**, The putative cholesterols between the inactive TM interface are shown. **b-e**, Phospholipid-like densities inside the TM bundles of inactive and active GABA_B_ structures. Fitting of the phosphatidylcholine, the most abundant phospholipid in mammalian cells, into the unassigned densities in inactive and active structures. The buried surface area of each phospholipid in inactive GB1(**a**), inactive GB2 (**b**), active GB1 (**c**) and active GB2 (**d**) is 1377 Å^2^, 1294 Å^2^, 1361 Å^2^, 1365 Å^2^, respectively. Inactive GB1 in blue; Inactive GB2 in yellow; Active GB1 in red; Active GB2 in green.

### Overall structure of GABA_B_ receptor in the inactive state

The overall architecture of GABA_B_ receptor reveals that heterodimeric GB1 and GB2 subunits face each other side-by-side with a pseudo C2-symmetry. The linker between extracellular VFT and TM domains within each subunit stands almost vertically to the bilayer. The extended ECL2 *β*-hairpin interacts extensively with the linker through three-stranded anti-parallel β-sheet, thus bridging VFT and TM domains in each subunit as a stalk (Fig. 1, 2a). This arrangement is similar to mGlu5 receptor^33^, yet lacking of cysteine-rich domain leading to the shorten height of the receptor and partially contributing to the enhanced stability of the receptor in the inactive conformation initially observed by negative stain EM (Extended Data Fig. 1d). The structure of the GABA_B_ VFT domains in the context of full-length receptor is very similar to that of the corresponding crystal structures of soluble VFT in both inactive and active states previously reported^29^, with root mean squared deviation (r.m.s.d.) of 0.8 Å and 1.2 Å, respectively (Extended Data Fig. 7a, b).

**Extended Data Fig.7.**
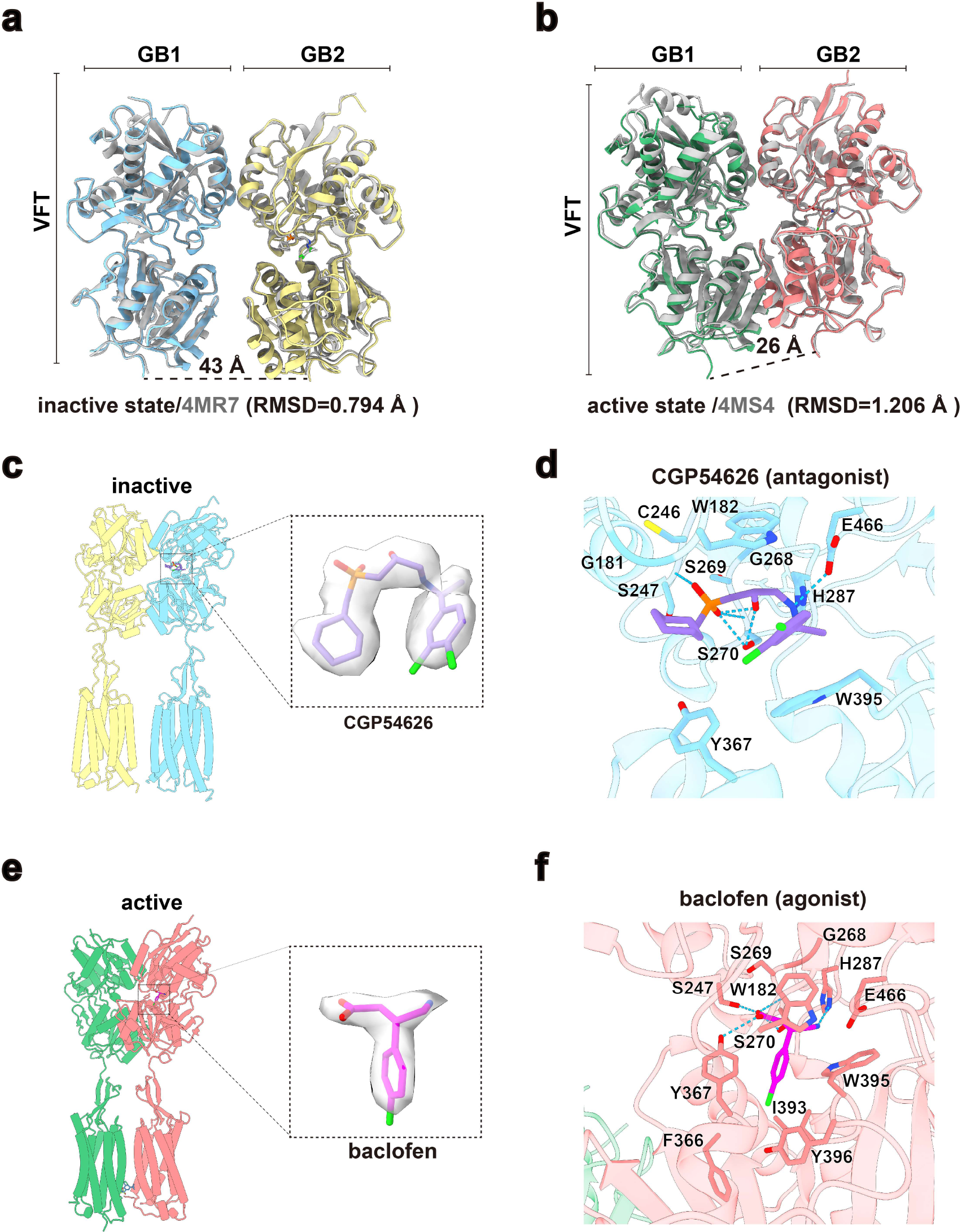
Structural comparisons of the VFT and ligand binding site. **a**, Superimposition of VFT domain from inactive structure of full-length GABA_B_ receptor and crystal structure of CGP54626-bound soluble VFT domain (grey; PDB code: 4MR7). **b**, Superimposition of VFT domain from active structure of full-length GABA_B_ receptor and crystal structure of baclofen-bound soluble VFT domain (grey; PDB code: 4MS4). **c-d**, Cryo-EM density (**c**) and the binding pocket analysis (**d**) of CGP54626 in inactive state. Detailed interaction of CGP54626 in the orthosteric binding site, in agreement with the reported crystal structure of CGP54626-bound GABA_B_ VFT. **e-f**, Cryo-EM density (**e**) and the binding pocket analysis (**f**) of baclofen in active state. Detailed interaction of baclofen in the orthosteric binding site is in agreement with the reported crystal structure of baclofen-bound GABA_B_ VFT.

CGP54626 anchors at the lobe1 of GB1 VFT by largely hydrogen bonds and maintain the receptor in the inactive state (Extended Data Fig. 7d). Accordingly, both VFT modules in the structure of the antagonist-bound receptor are in an open state, resulting in the heterodimer interfaces of GABA_B_ in the VFT region mainly through two lobe1s with the buried surface area of 702 Å^2^ (Extended Data Fig. 8a). In addition, the interfaces in the inactive state also involve the TM domains. Backbone separation distance of GABA_B_ between the TM5 helices are 8 Å in contrast to 17 Å of the inactive structure of mGlu5 (Extended Data Fig. 9a, b) and thus make extensive contacts via lipid-mediate Van der Waals interactions at the extracellular half and salt bridging at intracellular half (Fig. 3a-c). Salt bridge network close to the cytoplasmic membrane surface is mediated by a charged-residue quartet from the TM3 and TM5 helices, namely H689^3.55^ and E790^5.60^ of GB1, H579^3.55^ and E677^5.60^ of GB2 (superscript refers to the GPCRdb numbering scheme)^39^. Moreover, we observed well-resolved and ordered densities along the TM interface and assigned as putative cholesterols which connect a triplet of phenylalanine residues packing against one another in the proximal to the polar network (GB1a: F786^5.56^; GB2: F670^5.53^ and F673^5.56^) and a quintuplet of leucine residues in the middle of TM region (GB1a: L779^5.49^, L782^5.52^, L783^5.53^; GB2: L666^5.49^, L669^5.52^), all of which are from TM5 helices (Fig. 3b). At the extracellular half of the TM interface, TM5 helices facing each other with short uncharged sidechains form a deep crevice occupied by three putative cholesterols contributing to an additional buried surface area of 419 Å^2^, further substantially stabilizing the receptor in the inactive conformation (Fig. 3a, Extended Data Fig. 6a). This novel TM interface represents the structural signature of GABA_B_ heterodimer TM domains in the inactive conformation.

**Extended Data Fig.8.**
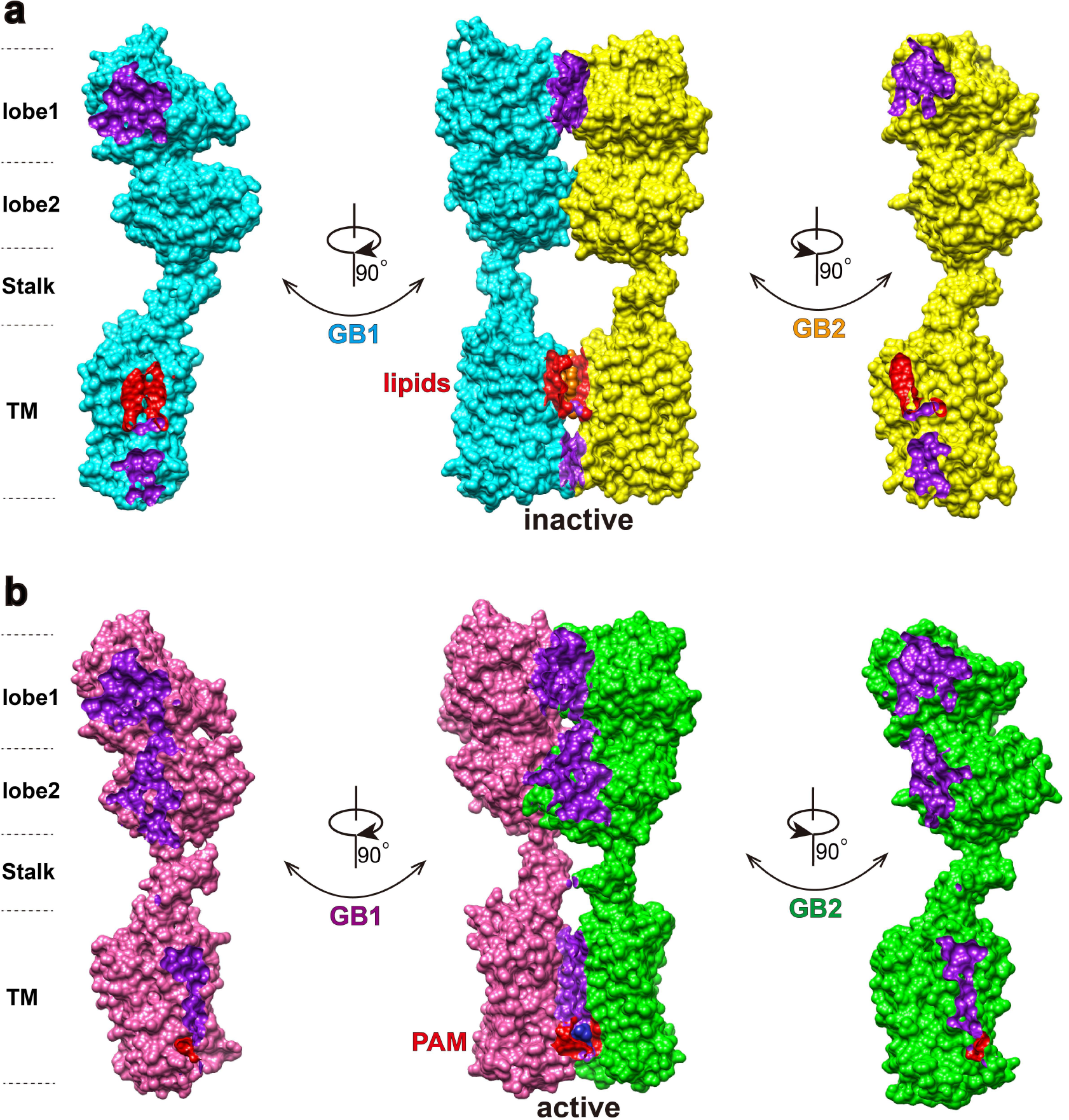
Comparison of the intersubunit interfaces in inactive and active GABA_B_ receptor. Intersubunit interfaces in inactive (**a**) and active (**b**) GABA_B_ receptor. The direct interface of GB1 and GB2 are colored purple. The putative cholesterols and the PAM interfaces are colored red.

**Extended Data Fig.9.**
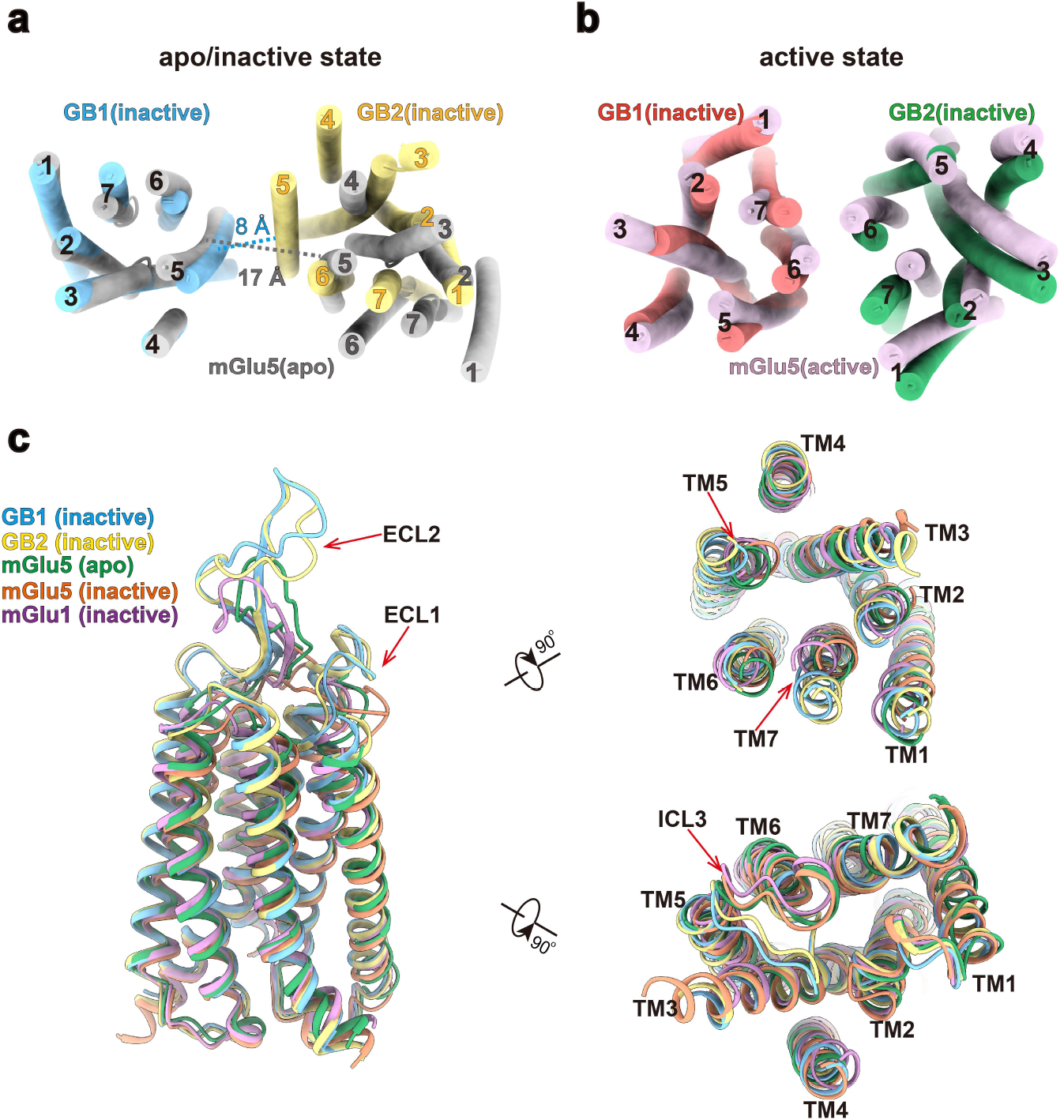
Structural comparison of GABA_B_ with mGlu receptors. **a-b**, Extracellular view of the superposed TM bundles of GABA_B_ and mGlu5 in inactive apo state (grey; PDB code: 6N52) (**a**) and active state (pink; PDB code: 6N51) (**b**). The TM5 helices of GABA_B_ receptor are closer compared with that of the inactive mGlu5 receptor measured at the closest separation of TM5 helices. **c**, Superimposition of TM domains of inactive GABA_B_ receptor, inactive mGlu1 (PDB code: 4or2) and inactive, apo mGlu5 (PDB code: 4oo9, 6N52). Red arrow represents the conformational difference of TM5, TM7, ECL1, ECL2, ICL3 between the GABA_B_ and the mGlu receptors.

### Conformational transition of GABA_B_ heterodimer during activation

Our 2.9 Å cryo-EM structure of the GABA_B_ receptor in the active state reveals distinct structural configuration in both VFT and TM domains to that in the inactive state (Fig. 2a). Agonist baclofen anchors between two lobes of the GB1 VFT (Extended Data Fig. 7f) and induces the closure of the GB1 VFT, with the lobe 2 of GB1 VFT undergoing a twist motion and contacting the opposite lobe2 shortening the distance between the C termini of the two VFT from 43 Å to 26 Å, in comparison to the distance changing from 45 Å to 32 Å as observed in the previous structural studies of the soluble VFT (Extended Data Fig. 7a, b). This conformational changes triggered by agonist binding propagates through the stalks and finally relays to the TM domains, leading to translations of the GB1 and GB2 TMs by 31 Å and 19 Å, respectively. Consequently, these structural transformations result in a TM6/TM6 interface (Fig. 2b, Fig. 3d), consistent with previous cross-linking studies^40^. This interface is also observed in the agonist-bound structure of the mGlu5 homodimer^33^, suggesting the TM6/TM6 interface is likely a hallmark of activation of class C GPCR.

Akin to the TM interface in the inactive GABA_B_ receptor, the interactions between the TM6/TM6 helices are strengthened by the hydrophobic and hydrophilic contacts. Compared to the lipid-mediated hydrophobic interactions at the extracellular half in the inactive state, the two TM6 helices in the active conformation directly pack together at their extracellular half via hydrophobic interactions, involving V814^6.48^, I818^6.52^, V822^6.56^, I825^6.59^, L826^ECL3^, A832^7.27^, F836^7.31^ from GB1 and M694^6.41^, I701^6.48^, M702^6.49^, I705^6.52^, A789^6.55^, V709^6.56^, L712^6.59^, T713^ECL3^, Q716^ECL3^, V719^7.27^ from GB2 and contributing to a buried surface area of 507 Å^2^ (Fig. 3d, Extended Data Fig. 8b). In the middle of the TM interface is a hydrogen bond network locked by the Y^6.44^ and N^6.45^ from both subunits, which are conserved through class C GPCR family (Fig. 3d, Extended Data Table 1). The TM6/TM6 interface is further stabilized by BHFF, a potent and selective GABA_B_ receptor PAM agonist (PAM-ago), which can enhance the potency of endogenous agonist GABA and also directly active GABA_B_ receptor in the absence of other agonists with the mechanism of act unknown^41^. The high-resolution cryo-EM map enables us identify and unambiguously assign the BHFF into the density located in the cavity formed by the intracellular tips of TM5-TM6 of GB1 and TM6 of GB2 (Fig. 3e, f). It’s noteworthy that we found the pure enantiomer (+)-BHFF fitting into the density better than (*−*)-BHFF, in agreement with previous pharmacological studies which suggested that rac- and (+)-BHFF could enhance the GABA potency by 15.3-fold and 87.3-fold, respectively. The (+)-BHFF is mainly recognized by the hydrophobic interactions, including A788^5.58^, Y789^5.59^, M807^6.41^, Y810^6.44^ of GB1 and K690^6.37^, Y691^6.38^, M694^6.41^ of GB2. Additionally, 3-hydroxy group and ketone are hydrogen-bonded to K792^ICL3^ of GB1(Fig. 3f). Given that BHFF can directly activate the GABA_B_ receptor and both its potency and efficacy could be enhanced in the presence of GABA, we anticipate that the cavity where BHFF occupied might be allosterically modulated by the orthosteric binding pocket and *vice versa*. We thus identified a novel PAM-binding pocket within a GPCR dimer and thus provide a template for future structure-based drug design of allosteric modulators targeting class C GPCR dimerization interface.

**Extended Data Table 1.**
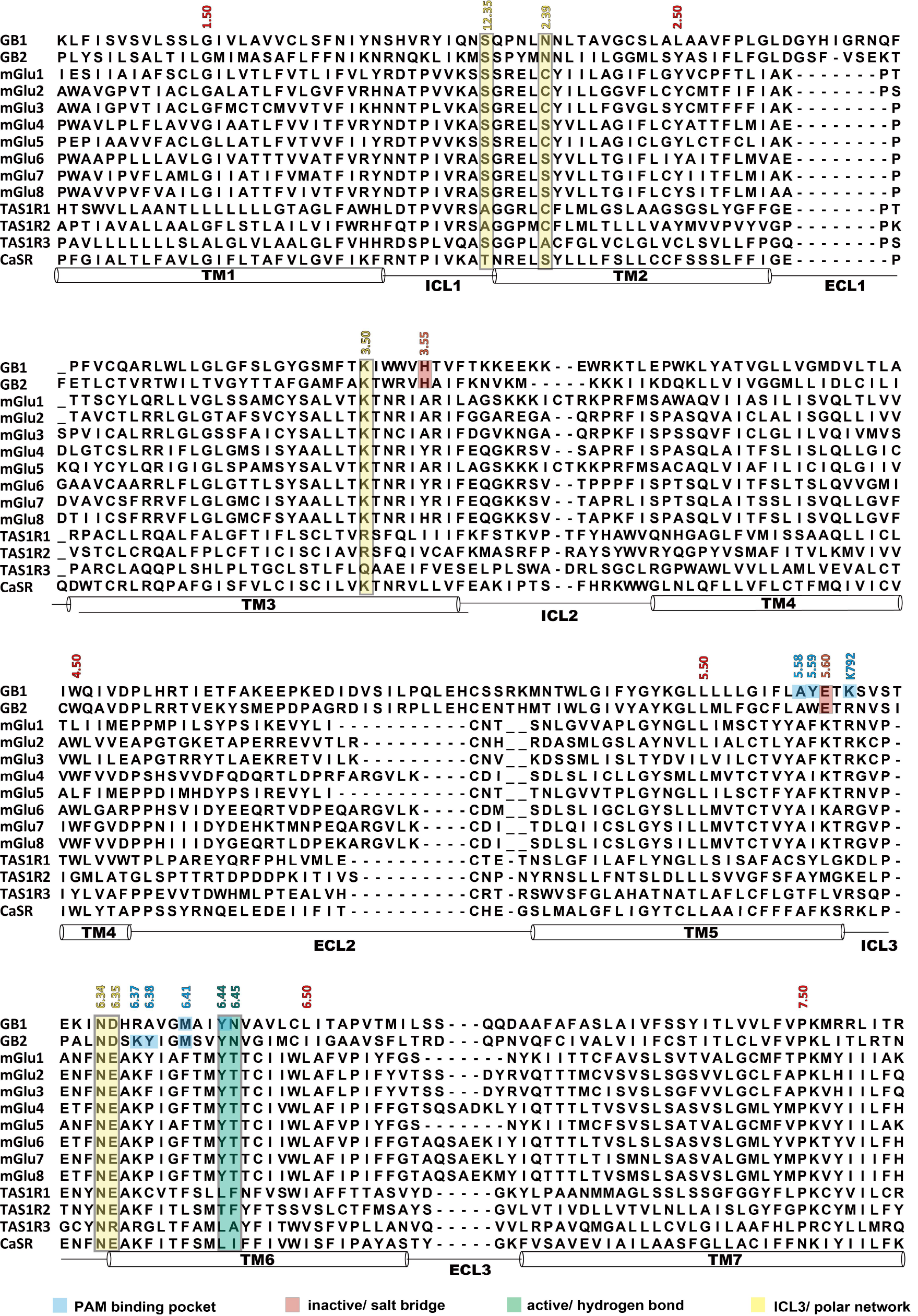
Sequence alignment of TMs of class C GPCRs. The sequence alignment of TMs of class C GPCRs was created using GPCRdb (http://www.gpcrdb.org) and Jalview software. Secondary structure elements are annotated underneath the sequences. Highlighted are residues involved in PAM binding pocket (blue), salt bridge in inactive TM interface, hydrogen bond in active TM interface (green) and polar network stabilizing ICL3 (yellow).

**Extended Data Table 2.** Cryo-EM data collection, refinement and validation statistics.

|  | Inactive GABA<sub>B</sub> receptor | Active GABA<sub>B</sub> receptor |
| --- | --- | --- |
| **Data collection and processing** |  |  |
| Magnification | 4,9310 | 4,9310 |
| Voltage (kV) | 300 | 300 |
| Electron exposure (e<sup>-</sup>/Å<sup>2</sup>) | 62 | 62 |
| Defocus range (μm) | -0.5 ~ -2.5 | -0.5 ~ -2.5 |
| Pixel size (Å) | 1.014 | 1.014 |
| Symmetry imposed | C1 | C1 |
| Initial particle images (no.) | 2,969,413 | 3,075,533 |
| Final particle images (no.) | 374,589 | 237,606 |
| Map resolution (Å) |  |  |
| (Overall/VFT/TM) | 2.9/2.8/3.0 | 3.0/2.9/3.0 |
| FSC threshold | 0.143 | 0.143 |
| Map resolution range (Å) | 2.2-4.0 | 2.2-4.0 |
| **Refinement** |  |  |
| Initial model used (PDB code) | 4MR7 | inactive GABA<sub>B</sub> structure |
| Model resolution (Å) | 2.9 | 3.3 |
| FSC threshold | 0.5 | 0.5 |
| Model resolution range (Å) | 2.5-4.0 | 2.5-4.0 |
| Map sharpening *B* factor (Å<sup>2</sup>) |  |  |
| (Overall/VFT/TM) | 83.6/84.1/67.6 | 78.5/80.9/190.3 |
| Model composition |  |  |
| Non-hydrogen atoms | 10915 | 10894 |
| Protein residues | 1355 | 1348 |
| Ligands | 9 | 10 |
| *B* factors (Å<sup>2</sup>) |  |  |
| Protein | 61.64 | 66.88 |
| Ligand | 63.76 | 88.08 |
| R.m.s. deviations |  |  |
| Bond lengths (Å) | 0.006 | 0.004 |
| Bond angles (°) | 0.741 | 0.684 |
| Validation |  |  |
| MolProbity score | 1.69 | 1.65 |
| Clashscore | 8.01 | 7.15 |
| Poor rotamers (%) | 0.00 | 0.00 |
| Ramachandran plot |  |  |
| Favored (%) | 96.27 | 96.18 |
| Allowed (%) | 3.73 | 3.82 |
| Disallowed (%) | 0.00 | 0.00 |

### Structure and activation of TM domain of GABA_B_ receptor

All the available TM structures of class C GPCRs are in the apo or inactive conformation including cryo-EM structure of the full-length mGlu5 bound to agonist^33^. Superposition of individual TM domains between GABA_B_ and other class C GPCRs including mGlu1 and mGlu5 reveals distinct structural features for TM5, TM7, ECL1, ECL2 and ICL3 (Extended Data Fig. 9c). Extracellular tips of TM5 and TM7 helices of GABA_B_ receptor demonstrate obvious outward shifting by 5 Å, likely resulting from the presence of endogenous phospholipid in the center of TM bundle which extensively contacts TM3, 5, 6 and 7 (Extended Data Fig. 6a). ECL2 of GABA_B_ spanning in a β-hairpin configuration is 5-residue extended relative to other class C GPCRs, leading to more tight interaction with the linker and enabling the direct contacts with the bottom of VFT (Fig. 4, Extended Data Fig. 9c). Therefore, this extension facilitates the conformational changes of VFT induced by agonist binding conducting to TM domain. Moving to the intracellular loops, the most profound difference is the position of ICL3, which is proposed to play a crucial role in G-protein coupling^25^. The ICL3 of GABA_B_ receptor is comprised of 11 residues for both subunits (residues 791-801 for GB1 and residues 678-688 for GB2) and stretched over the TM pocket at the cytoplasmic surface. The conformation of ICL3 is stabilized by the polar network formed by S515^12.53^, N520^2.39^, K574^3.50^, N687^6.34^ and D688^6.35^ which are highly conserved among class C GPCRs and further strengthened by hydrophobic interactions between L511^12.49^, M514^12.52^, A675^5.58^, W676^5.59^ (Extended Data Fig. 10a-c). We anticipate that the unique conformation of ICL3 in GABA_B_ receptor restricts the access of G proteins or β-arrestins to the receptor similar to the role of helix 8 of the angiotensin II receptor AT_2_R^42^. In contrast, the same loop from mGlu1 and mGlu5 is 2-residue shorter and stands almost perpendicular to the bilayer, leaving the TM cavity at the cytoplasmic side more accessible to the effector proteins (Extended Data Fig. 9c).

**Fig.4.**
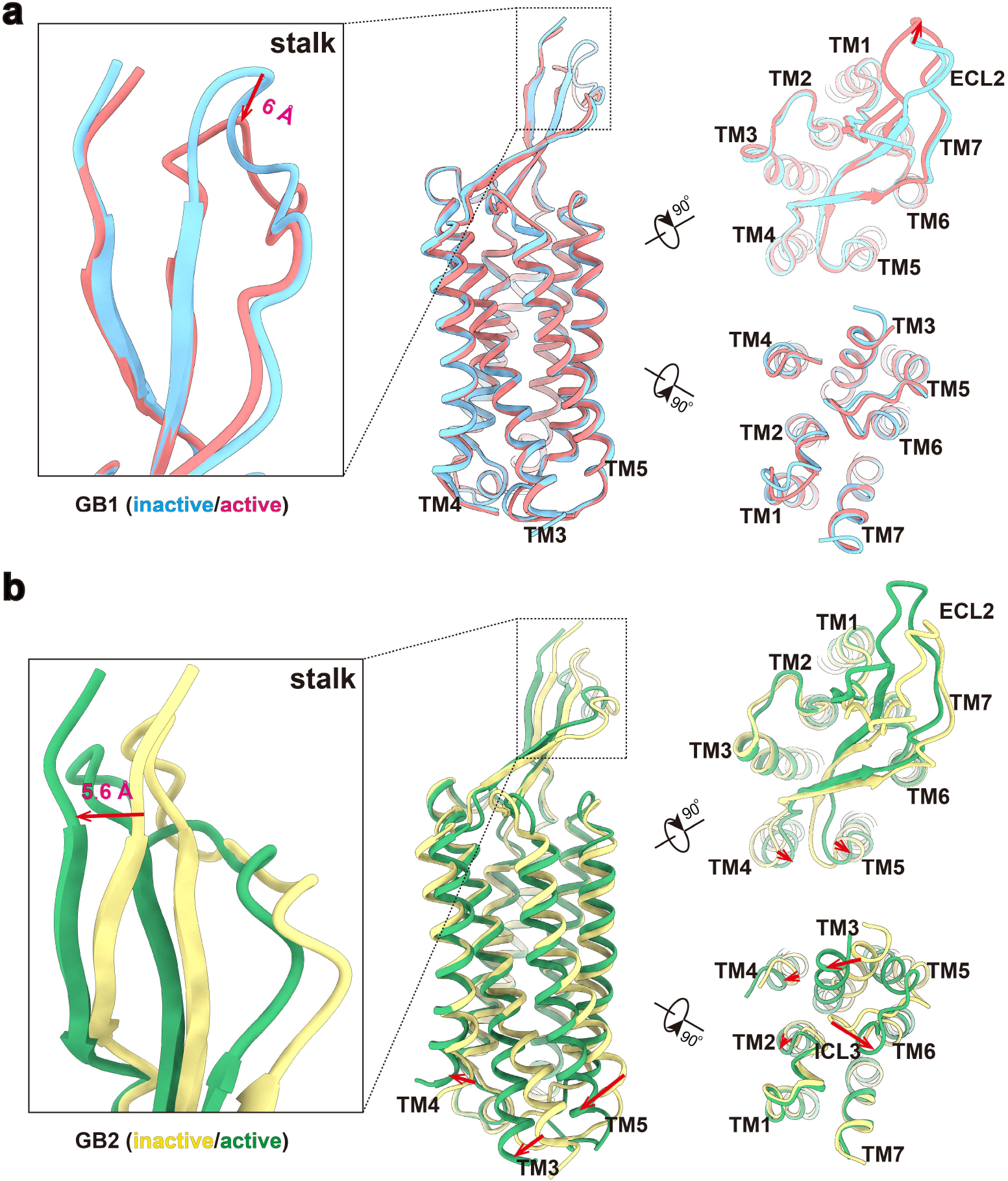
Structural comparison of individual TM domains between inactive and active states. GB1 TM domain (**a**) and GB2 TM domain (**b**) between inactive and active states are superposed respectively. Side, extracellular and intracellular views are shown. Magnified views of the stalk domains are shown in the left. Red arrows represent the movement direction and distance of TMs and loops.

**Extended Data Fig.10.**
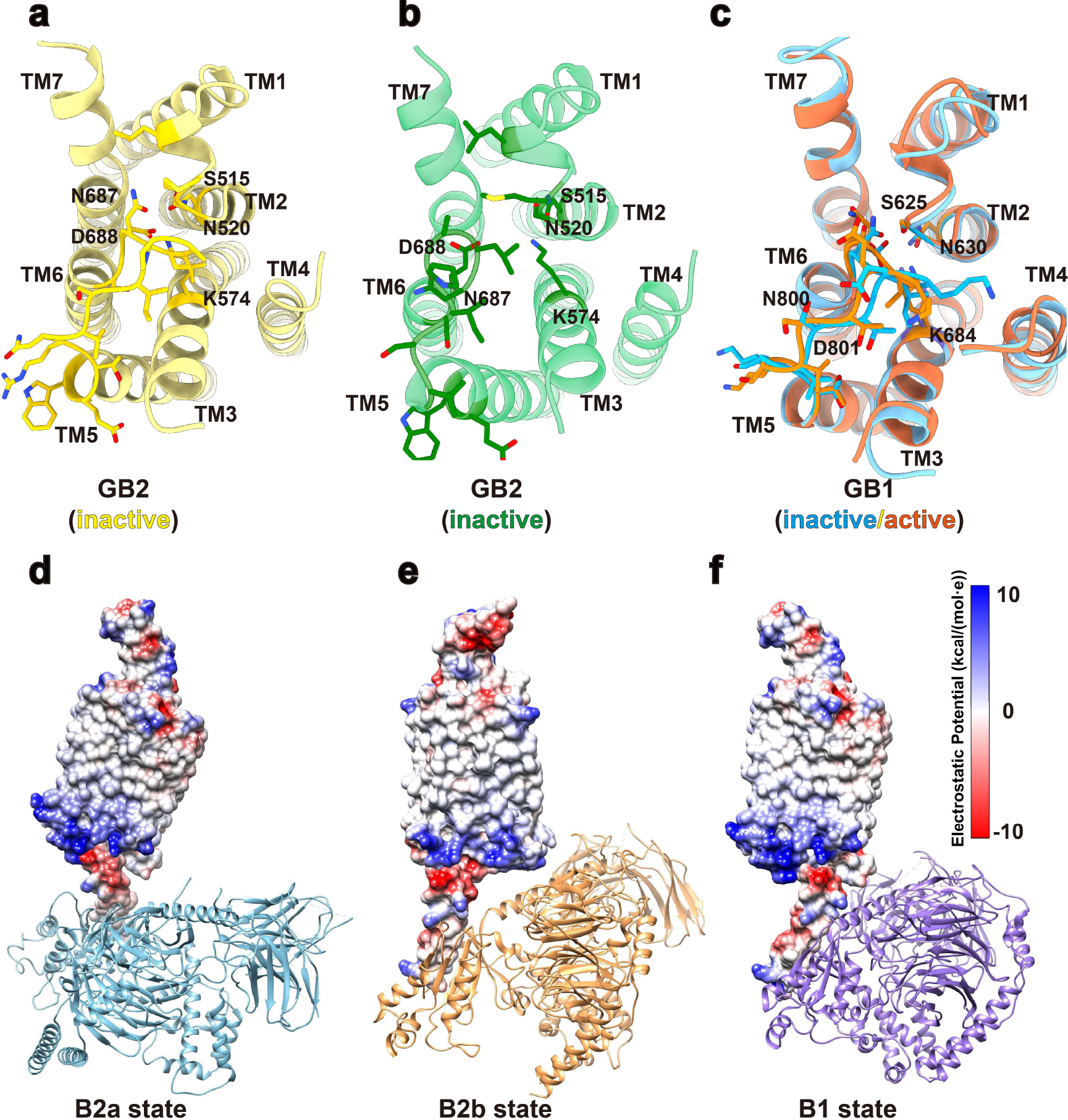
Conformation of ICL3 in inactive and active GABA_B_ structures and the α5–receptor interface in GABA_B_–G_il_ complex. **a-c**, Interactions of the ICL3 with other residues of TM domain in GB2 inactive (**a**), active state(**b**) and GB1 in both states(**c**). **d-f**, Electrostatic charge surface illustrating the interface of with the GABA_B_ receptor in B2a (**d**), B2b (**e**)and B1(**f**) states, showing the complementary-charge distribution between the C terminus of the α5-helix and the cytoplasmic surface of TM bundle.

The overall structures of the GABA_B_ TM domains are highly similar in the inactive state with r.m.s.d. of 0.939 Å as expected, considering that the two TM domains share 72% sequence homology. Although upon activation the apical tip of GB1 ECL2 shifts 6 Å measured at Cα atom of I750^ECL2^ residue, the extracellular conformational changes induced by agonist binding do not relay to GB1 TM domain through the stalk, which is mostly unchanged (r.m.s.d. of 2.23Å) (Fig. 4a). In contrast, substantial conformational changes occur at GB2 stalk region (r.m.s.d. of 3.94 Å) translating by 5.6 Å measured at I469, triggering the extracellular tips of TM4 and TM5 shifting 2.0 Å and 2.9 Å to the opposite direction measured at D619^4.54^ and T654^5.37^, respectively (Fig. 4b). Consequently, the intracellular tip of GB2 TM3 undergoes 5.2 Å shifting (measured at Cα atom of I581^3.57^) towards TM4 and TM2, leading to a 2.7 Å and 1.7 Å outward movement measured at Cα atom of L598^4.33^ and at Cα atom of M519^2.38^, respectively. The most profound conformational changes at the cytoplasmic half locate at TM5-ICL3. GB2 TM5 extends one helical turn and shifts 5.4 Å measured at Cα atom of W678^5.61^ toward TM3, resulting in the breaking of the ionic interactions between N687^6.34^, D688^6.35^and S515^12.53^, N520^2.39^, K574^3.50^, and thus the ICL3 stands up uncovering the lid of TM pocket at the cytoplasmic side (Extended Data Fig. 10b). Taken together, the orchestration of the conformational changes in the TM domain of GB2 subunit enables it accommodating the down-streaming Gi/o protein to initiate the intracellular signaling.

### Gi heterogeneously coupling to GABA_B_ receptor heterodimer

The structural rearrangement of GB2 TM domain barely exposes residues from TM helices and thus creates a shallow binding site for the heterotrimeric G_i1_ protein (Fig. 5a-c), whereas all the available GPCR–G structures show a deep cavity formed by the outward movement of the cytoplasmic half of TM6 together with TM2, 3, 5, and 7^43^. This remarkable shallow pocket disallows the insertion of the α5-helix of the Gαi Ras-like domain into the receptor TM domain and is probably responsible for the conformational heterogeneity present in the GABA_B_–G_i1_ complex. We identified three thermostable conformations demonstrating well-resolved density for G_i1_ protein, which are B2a, B2b and B1 states with the population distribution of 74%, 9% and 17% among the sub dataset, respectively (Fig. 5a-c, Extended Data Fig. 3), this observation suggests that G_i1_ protein predominantly couples to GB2 subunit which is consistent with the asymmetric activation model of GABA_B_ receptor. Superposing the isolated GB1–G_i_ structure from B1 state to the heterodimer structure of either B2a or B2b state shows the potential steric hindrance between the two Ras-like domains of G_i1_, perfectly explaining why only one G protein binds to the GABA_B_ receptor at a time (Fig. 5d).

**Fig.5.**
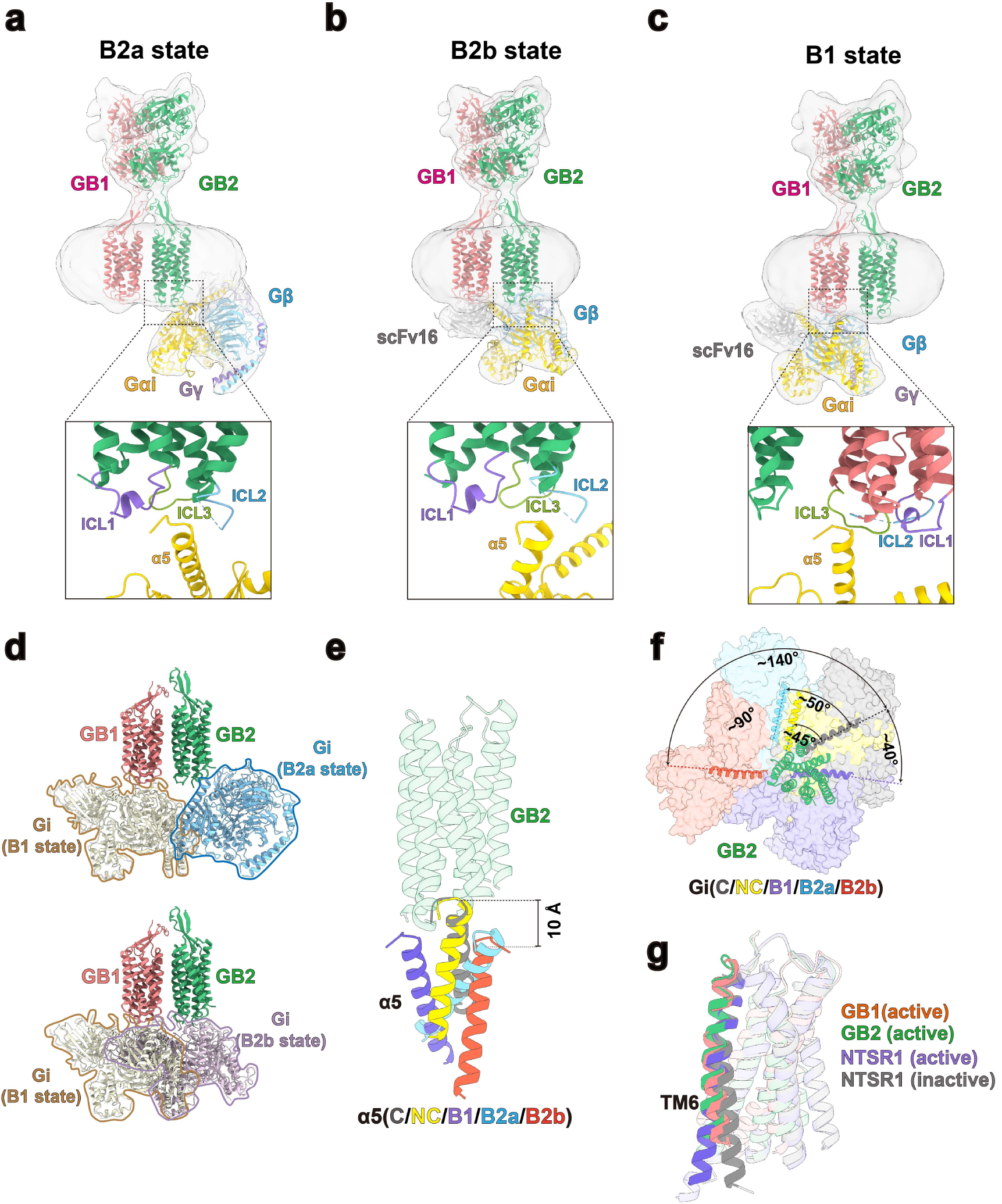
Three distinct modes of Gi coupling to GABA_B_ receptor. **a-c**, Cryo-EM density maps and the models of the GABA_B_–G_i1_ complex in B2a state (**a**), B2b state (**b**), and B1 state (**c**). Magnified views of the interaction between α5-helix of Gαi and GABA_B_ receptor are shown in the bottom. GB1 in red, GB2 in green, Gαi in gold, Gβ in cyan, Gγ in purple and scFv16 in grey. Intracellular loops 1, 2, and 3 are highlighted. **d**, Superposition of the B1 state with either B2a state or B2b state showing the potential steric clash between Gi proteins, indicating that only one Gi protein can bind to GABA_B_ receptor at one time. Gi in B2a state, brown; Gi in B2b state, blue; Gi in B1 state, grey. **e**, Comparison of the Gα_i1_ α5-helix binding position between GB1 (B1 state), GB2 (B2a and GB2b states) and the NTSR1– Gi1 protein complex in canonical (C; PDB code: 6OS9) and noncanonical (NC; PDB code: 6OSA) states. The position of α5-helix from GABA_B_–Gi complex in different states show around 10 Å downward movement compared with that of NTSR1–G_i1_ complex. Structures were aligned by the TM domains and only GB2 TM domain is shown for clarity. Color usage for α5: C state, grey; NC state, gold; B1 state, purple; B2a state, cyan; and B2b state, red. **f**, Orientations of the Gi protein when coupling to GABA_B_ and NTSR1 receptors. Rotation of Gi protein are measured by the αN domains. G proteins are coloured as described in **e. g**, Comparison of the active GB1 and GB2 TM domains with the inactive and active structures of NTSR1, showing the positions of TM6 helices of GB1 and GB2 subunits are similar to that of NTSR1 in inactive state.

The interface between the activated GABA_B_ receptor and G_i1_ protein in B2a state involves ICLs and intracellular tip of TM3 of the receptor and α5-helix of the Gαi Ras-like domain, primarily stabilized by hydrophobic interactions and complementary change interactions (Fig. 5a, Extended Data Fig. 10d). The carboxyl terminal α5-helix of the Gα_i_ overlaps the position of GB2 ICL3 in the inactive state, explaining the requirement for GB2 ICL3 opening to accommodate the G_i/o_ protein. Previous studies showed that the single mutation L686P in the ICL3 of GB2 suppresses the activation of G protein in either HEK293 cells or cultured neurons, highlighting the critical role of ICL3 of GB2 for the coupling of the heterodimeric GABA_B_ receptor to G-proteins^25^, in agreement with our structural findings. Consistently, the α5-helix is also the sole coupling site with the receptor in the other two states (Fig. 5b, c, Extended Data Fig. 10e, f). Interestingly, the α5-helix of the Gαi in three GABA_B_–G_i1_ states translates downwards to the similar extent (about 10 Å) relative to that of class A GPCR–G_i_ complex in the canonical and noncanonical states (Fig. 5e), leading to the absence of Gβγ in the receptor– G_i1_ interface. However, G_i1_ in B2b and B1 states is rotated by about ∼90° to opposite side of that in B2a state relative to the receptor, positioning G_i1_ in the complex in distinct arrangement (Fig. 5f). Surprisingly, unlike class A and B GPCRs, TM6 holds its position in response to activation, constrained by the TM6/TM6 interface in the active state (Fig. 5g).

Intriguingly, the observation of the GABA_B_–G_i1_ complex in GB1 state provides the structural explanation of the basal activity of GABA_B_ receptor given that GB1 TM domain shares the same conformation in the inactive and active states, in agreement with that expression of GB1 alone at the cell surface activates ERK1/2 through G_i/o_-dependent pathway^27,28^. Considering the significant difference at population between the two B2 states, we anticipate that B2a is likely more thermodynamically stable and therefore forms after the B2b state along the activation pathway of G protein. Similar phenomenon was observed in the neurotensin receptor 1(NTSR1)–G_i1_ complex^44^.

## Conclusion

GABA_B_ receptor represents the first example of an obligatory GPCR heterodimer, functions as an allosteric multi-domain protein and oscillates between the inactive and active states^21^. Ligands or G proteins interplay with the pre-existing states of the receptor based on “selection mode of allostery”^21,45^. In this study, by synchronizing receptor in the inactive state with the presence of antagonist or completely stabilizing receptor in the active state with the presence of agonist, PAM and G_i1_ protein, we determined the atomic-resolution cryo-EM structures of a human full-length GABA_B_ heterodimer in both inactive and active states. The high-resolution cryo-EM maps not only confirmed the binding mode of CGP54626 and baclofen observed in the crystal structures of GABA_B_ VFT^29^, but also revealed the working mechanism of BHFF as PAM agonist, which stabilizes a novel intersubunit interface in the TM domains of the active GABA_B_ receptor. Surprisingly, endogenous phospholipids settle inside each TM extracellular pore, equivalent to classical orthosteric binding pocket for class A GPCR, in all GABA_B_ structures obtained in this work and lead to a distinct conformational configuration of GABA_B_ TM domains compared to other class C GPCRs, suggesting the presence of these lipids may function as structural components which is probably a unique phenomenon to GABA_B_ receptor.

Structural comparison of GABA_B_ receptor in different states enabled us to propose a structural rearrangement for GABA_B_ receptor activation. Agonist binding induces the compaction of two VFTs (Fig. 2, Extended Data Fig. 7a, b), propagating through the stalk domains to the TM domains, which in turn reorients the TM interface between two subunits from TM3-TM5/TM3-TM5 in the inactive state to TM6/TM6 in the active state, in perfect agreement with previous crosslinking data for GABA_B_ receptor and mGlu homodimers^40,46^. Therefore, the overall domain rearrangements upon activation of GABA_B_ receptor resemble the published cryo-EM structures of mGlu5^33^, suggesting this reorientation of TM domains is possibly the hallmark of class C GPCR activation.

We determined cryo-EM structures of a GPCR heterodimer in a complex with G_i1_ protein, which only can couple to either one of the subunit at one time to avoid steric hindrance by the presence of Ras-like domain of two Gα proteins. The unprecedented structures reveal novel conformational changes within the GB2 TM domain to accommodate the G_i_ protein involving the opening of ICL3 and movement of TM3, 4 and 5 helices, enabling GABA_B_ receptor predominantly couples to G_i1_ protein through GB2 subunit. However, the structure of GABA_B_– G_i1_ complex coupling via GB1 demonstrates that GB1 couples to G_i_ protein at certain circumstance. Indeed, GB1 in C. elegans directly couples to G_i/o_ protein-dependent signaling^47,48^, indicating that GB1 retains the capacity for Gi protein coupling during evolution. Common activation mechanism for G protein activation by GPCRs has been proposed, in which the landmark of GPCR activation is the outwards movement of TM6 forming a cavity to accommodate the G proteins or β-arrestins, reflecting the convergence of activation pathways in class A, B and F GPCRs^43,49,50^. However, the GABA_B_–G_i1_ engagement patterns have not been observed in the structures of other GPCR-G complexes, implying the complex of GPCR-G complex, especially for GPCR in dimeric form. Further studies are required to determine the high-resolution structure of GABA_B_–G_i1_ complex to decipher the molecular mechanism of GABA_B_–G protein assembly. Additional structural studies are also needed to determine whether similar G protein engagement are present in other class C GPCRs.

## Acknowledgements

We thank S. Chang for technical support in cryo-EM data collection at the Center of Cryo-Electron Microscopy, Zhejiang University; C. Ma for support in protein purification at the Protein Facilities, Zhejiang University School of Medicine. J.L. was supported by the Ministry of Science and Technology (2018YFA0507003), the National Natural Science Foundation of China (NSFC) (81720108031, 81872945, 31721002 and 31420103909), the Program for Introducing Talents of Discipline to the Universities of the Ministry of Education (B08029), and the Mérieux Research Grants Program of the Institut Mérieux. Y.Z. was supported by the National Science Foundation of China (81922071) and the National Key Basic Research Program of China (2019YFA0508800).

## Author contributions

C.S. designed the constructs, expressed and purified the antagonist-bound GABA_B_ and the agonist/PAM stimulated GABA_B_-G_i1_ complex for cryo-EM data collection with the help of S.Z.; C.S., R.Z. and C.L. performed pull-down assay; C.L. and L.-N.C. expressed and purified scFv16; C.X. performed BRET assay; D.-D.S. evaluated the sample by negative-stain EM; C.M. prepared the cryo-EM grids; C.M., D.-D.S. collected the cryo-EM data; C.M. performed cryo-EM map calculation, model building and structure refinement; C.M. and C.S. analyzed the structures and prepared the figures with the help of D.-D.S., Q.S. and Z.J.; Y.Z. and J.L. conceived and supervised the project, analyzed the structures, and wrote the manuscript with input from all the authors.

## Methods

### Constructs

Human GABA_B_ receptor with the HA signal peptide including GB1a (UniProt: Q9UBS5) and GB2 (UniProt: O75899) were cloned into pEG BacMam vector, respectively^51^. To facilitate expression and purification, an 8× histidine tag and 3C protease cleavage site were inserted at the C-terminus of GB1a subunit, while a Flag epitope tag (DYKDDDD) and a 2×GSG linker were added to the N-terminus of GB2 subunit. Both GB1 and GB2 with a long, flexible C terminal affected the level of expression. Therefore, we utilized different constructs with cleavable C-terminal domains: GB1 and GB2 were truncated after coiled-coil domain; GB1△cc and GB2△cc were truncated before coiled-coil domain; GB1△C means the C-terminus of GB1 was removed. In addition, we construct the Dual pEG BacMam vector could express GB1 and GB2 by combining two plasmids using homologous recombinant enzyme to increase expression level.

### Expression and purification of inactive GABA_B_ receptor

GB1a (GB1cc residues 15-919) and GB2 (GB2cc residues 42-819) were co-expression in HEK293F cells. In brief, purified plasmid DNA was mixed with PEI 25K in a 3:1 ratio of PEI to DNA (w/w) followed by addition to HEK293F cells when density reached around 2.8 million/ml^52^. 10 mM sodium butyrate was added after 16-18 hours post-infection, then cells grown for 3 days at 30 °C before harvested^51,52^. The infected cells were collected by centrifugation at 1000 g for 15 min and washed once with 1×PBS buffer. Cells were suspended in 50 mM HEPES pH 7.5, 150 mM NaCl, 10% glycerol, 2 mM MgCl_2_, 20 µM CGP54626 (Tocris Bioscience) with cocktail followed by homogenization. The membrane was solubilized for 3 hours at 4 °C with 0.5% (w/v) lauryl maltose neopentyl glycol (LMNG, Anatrace), 0.1% (w/v) cholesteryl hemisuccinate (CHS, Anatrace). After centrifugation at 30,000 g for 30 min, the supernatant was bound to Ni-NTA column and further loaded onto M1 anti-FLAG affinity resin. The protein was eluted in elution buffer consisting of 50 mM Hepes pH 7.5, 150 mM NaCl, 0.01% LMNG, 0.002% CHS, 20 µM CGP54626, 5 mM EGTA and 0.1 mg/mL FLAG peptide. The GABA_B_ receptor was concentrated in a 100-kDa cutoff Vivaspin (Millipore) filter and run on a Superose™ 6 Increase column (GE Healthcare).

### Expression of heterotrimeric Gi

Heterotrimeric Gi was expressed as previously described^53^. In general, the dominant-negative rat Gαi1 was introduced four mutations (S47N, G203A, E245A and A326S) and a 6× histidine tag was added at the N-terminus of the β subunit. The Sf9 insect cells (Expression Systems) at a density of 2.4 million/ml were infected with both Gαi and Gβγ virus in a 1:1 ratio. Cells were harvested 48 hours after infection and collected by centrifugation at 1000g for 15 min. Finally, the cells were washed once with 1×PBS buffer and snap frozen in liquid nitrogen for later use.

### Expression and purification of scFv16

The secreted scFv16 was expressed and purified by using the bac-to-bac system as previously described^36^. In brief, the virus of scFv16 with 6× histidine tag at the C-terminus was infected in Trichoplusia ni Hi5 insect cells for 48 hours. The pH of Supernatant was balanced by addition of Tris pH 8.0 while Chelating agents were removed by addition of 1 mM nickel chloride and 5 mM calcium chloride. After incubation with stirring at room temperature for 1 hour the supernatant was loaded onto Ni-NTA resin and further eluted in elution buffer consisting of 20 mM Hepes pH 7.5, 500 mM NaCl, and 250 mM imidazole. The sample was first treated with 3C protease, then diluted and reloaded onto the Ni-NTA column to remove the 6×histidine tag. The flow-through was collected and purified over gel filtration chromatography using a Superdex 200 column. Finally, the concentrated of scFv16 was flash frozen in liquid nitrogen until further use.

### Formation and purification of the GABA_B_-G_i_-scFv16 complex

GB1a (GB1△C residues 15-860) and GB2 (GB2△cc residues 42-780) were chosen to form GABA_B_-G_i1_ complex. The conditions for GABA_B_ expression in HEK293F cells described earlier. Purified plasmid DNA was mixed with PEI 25K in a 3:1 ratio of PEI to DNA (w/w) was added to HEK293F when density reached around 2.8 million/ml. The infected HEK293F cells and the expressed G_i1_ cells together were suspended and disrupted in 50mM HEPES pH 7.5, 150 mM NaCl, 10% glycerol, 2 mM MgCl_2_, 100 µM baclofen and 50 µM BHFF (Tocris Bioscience) with cocktail followed by addition of 50 mU/ml apyrase and 1.0 mg scFv16. The membrane was solubilized with 0.5% (w/v) LMNG and 0.1% (w/v) CHS for 3 hours at 4 °C and separated by centrifugation at 30,000g for 30 min. The supernatant was bound to Ni-NTA column and further loaded onto M1 anti-FLAG affinity resin. The protein was eluted in elution buffer consisting of 50 mM Hepes pH 7.5, 150 mM NaCl, 0.01% LMNG, 0.002% CHS, 100 µM baclofen and 50 µM BHFF, 5 mM EGTA and 0.1 mg/mL FLAG peptide. The GABA_B_-Gi-SCFV16 complex was concentrated in a 100-kDa cutoff Vivaspin (Millipore) filter and run on a Superose™ 6 Increase column.

### G protein pull-down analysis

Three different group truncations of GABA_B_ receptor were cloned into pEG BacMam vector, respectively. Flag epitope tag (DYKDDDD) and a 2×GSG linker were added to the N terminus of GB2. Different group truncations of GABA_B_ receptor were expression in 30 ml HEK293F cells while Heterotrimeric Gi was expression in 30 ml Sf9 insect cells. These cells were suspended in 50 mM HEPES pH 7.5, 150 mM NaCl, 10% glycerol, 2 mM MgCl_2_, 100 µM baclofen and 50 µM BHFF with cocktail followed by homogenization. 50 mU/ml apyrase and 0.03mg scFv16 were added to incubate 3 hours. The membrane was solubilized with 0.5% (w/v) L-MNG and 0.1% (w/v) CHS for another 3 hours at 4 °C. After centrifugation, the supernatant was incubated with 25µl M1 anti-FLAG affinity resin for 1 hour at 4 °C and then eluted in 50 µl elution buffer supplemented with 100 µM baclofen and 50 µM rac-BHFF, 5 mM EGTA and 0.1 mg/mL FLAG peptide. Sample collected from different truncations were analyzed by SDS-PAGE.

### Bioluminescence resonance energy transfer (BRET) assay

HEK293 cells were transfected with wild type or truncation (GB1△C-GB2△cc) GABA_B_ receptor, Gβ1, Venus-Gγ2 and Gαi-Rluc8 by lipofectamine 2000 and split into 96-well flat-bottomed white microplates. After 24 h transfection, cells were washed and starved in PBS at 37 °C for 1 hour. BRET measurements were performed using the Mithras LB 940 (Berthold Technologies, German). The signals emitted by the donor (460–500 nm band-pass filter, Em 480) and the acceptor entity (510–550 nm band-pass filter, Em 530) were recorded after the addition of 5 µM Coelenterazine H. All measurements were performed at 37 °C. The BRET signal was determined by calculating the ratio of the Em 530 and Em 480. The net BRET ratio was defined as the experimental BRET signal values with the baseline subtracted (basal BRET ratio), which was recorded before the stimulation of cells. For dose-response experiments, data were analyzed using nonlinear curve fitting for the log (agonist) vs. Response (three parameters) curves in GraphPad Prism software.

### Cryo-EM grid preparation and data collection

For the preparation of Cryo-EM grids, 3 µl of the purified antagonist bound GABA_B_ heterodimer or baclofen/BHFF bound–GABA_B_–G_i1_ complex at ∼2.0 mg/ml was applied onto a glow-discharged 200 mesh holey carbon grid (Quantifoil R1.2/1.3). The grids were blotted and then plunge-frozen in liquid ethane using Vitrobot Mark IV (Thermo Fischer Scientific). Cryo-EM data collection was performed on a Titan Krios at 300 kV accelerating voltage in the Center of Cryo-Electron Microscopy, Zhejiang University (Hangzhou, China). Micrographs were recorded using a Gatan K2 Summit direct electron detector in counting mode with a nominal magnification of 29,000 ×, corresponding to a pixel size of 1.014 Å. Image stacks was obtained at a dose rate of about 8.0 electrons per Å^2^ per second with a defocus ranging from −0.5 to −2.5 µm. The total exposure time was 8 s and intermediate frames were recorded in 0.2 s intervals, resulting in an accumulated dose of 64 electrons per Å2 and a total of 40 frames per micrograph. A total of 4740 and 4624 movies were collected for the antagonist bound GABA_B_ heterodimer and baclofen/BHFF bound–GABA_B_–G_i1_ complex, respectively.

### Imaging processing and 3D reconstruction

For the dataset of antagonist-bound GABA_B_ heterodimer, image stacks were subjected to beam-induced motion correction using MotionCor2.1^54^. Contrast transfer function (CTF) parameters for each non-dose weighted micrograph were determined by Gctf^55^. Semi-automated particle selection were performed using RELION-3.0-beta2^56^, yielded 2,969,413 particles. The particles were extracted on a binned dataset with a pixel size of 2.028 Å and imported to CryoSPARC v2.42^57^ for 2 rounds of 2D classification. The poorly defined 2D classes were discarded, producing 1,757,222 particles for further Ab-initio reconstruction and heterogeneous refinement. After 2 rounds of heterogeneous refinement, the well-defined subset with 802,381 particles were re-extracted with a pixel size of 1.014 Å in Relion. The particles were subsequently subjected to 3D classification, produced two good subsets accounting for 610,689 particles. Further 3D classifications focusing the alignment on the complex and the TMD domains, produced high-quality subsets accounting for 374,595 particles, which were subsequently subjected to 3D refinement and Bayesian polishing. The overall refinement generated a map with an indicated global resolution of 3.0 Å at a Fourier shell correlation of 0.143. To further improve the map quality, local refinement focusing on the TMD and VFT domains were performed in Relion. The locally refined map for the VFT and TMD show a global resolution of 2.8 and 3.0 Å, which were merged using vop maximum command in UCSF chimera^58^. This composite map was then ‘Z-flip’ to get the correct handedness and used for subsequent model building and analysis.

For the dataset of GABA_B_-G_i1_ complex, movies were subjected to beam-induced motion correction using MotionCor2.1^54^ and CTF estimation using Gctf^55^. Template-based automated particle selection were performed using RELION-3.0-beta2, producing 3,075,533 particles for further 2D classification in CryoSPARC. The well-defined classes with 1,471,649 particles were selected for Ab-initio reconstruction and heterogeneous refinement, resulting in 488,801 particles with good density of the receptor. The subset was further re-extracted on a unbinned dataset in Relion and performed multiple rounds of 3D classification. Three classes showed different conformation of G protein binding (B1, B2a and B2b) were identified accounting for 30,542 particles, 130,901 particles and 16,048 particles, respectively. Whereas, other three classes showed the fuzzy density of the G protein. Subsets for the 3 conformational states were further subjected to 3D refinement and Bayesian polishing, yielded density maps with indicated global resolutions of 8.8 Å, 6.8 Å, 8.6 Å, respectively. To find out the TMD conformations of the G protein bound GABA_B_, the map of dominant conformation, B2a, was further refined with a mask on the receptor. The locally refined density map show a relatively clear density for the TM bundle arrangement with an indicated resolution of 5.8 Å. Due to the highly consistency of the receptor conformations in all 3D classes including three identified G protein-bound states and three poorly defined G protein-bound classes, we reasoned that the receptor adopts the same or highly similar conformation regardless of the orientation of G protein engagement. We then focused the alignment on the receptor and performed 3D classification with a mask on the receptor and TMD, respectively. The subsets accounting for 237,606 particles showed the high-quality EM density were selected and were subjected to 3D refinement and Bayesian polishing. The refinement generated a map with an indicated global resolution of 3.1 Å. To improve the quality of TMD domain, local refinement focusing on the TMD and VFT domains were further performed in Relion. The locally refined map for the VFT and TMD show a global resolution of 2.8 and 3.2 Å, which were merged in UCSF chimera and used for subsequent model building and analysis. Local resolution was determined using the Bsoft package^59^ with half maps as input maps.

### Model building and refinement

The initial template of the TMDs in GB1 and GB2 was generated using SWISS-MODEL^60^. The initial template of VFT domain was derived from the crystal structure of antagonist-bound GABA_B_ VFT (PDB 4MR7)^29^. Antagonist, agonist and PAM coordinates and geometry restraints were generated using phenix.elbow^61^. Models of TMD and VFT domains were docked into the EM density map of antagonist bound GABA_B_ using UCSF Chimera^58^. The initial model was subjected to iterative manual rebuilding in COOT^62^ and real-space refinement in PHENIX^61^. The final refinement statistics were validated using the module ‘comprehensive validation (cryo-EM)’ in PHENIX. The refined model of antagonist-bound GABA_B_ was used as the initial model for baclofen/BHFF-bound GABA_B_. Structures of TMD and VFT lobe domains of GB1 and GB2 were docked into the density map using UCSF Chimera, respectively. The model was further subjected to manual rebuilding and real-space refinement. The structures of baclofen/BHFF-bound GABA_B_ and G_il_ protein from the structure of human cannabinoid receptor 2-G_i1_ complex (PDB 6PT0)^63^ were used to generate the docked model of GABA_B_-G_i_ complex in B1, B2a and B2b states, respectively. The refinement statistics of the inactive and active GABA_B_ heterodimer are provided in Table S1. Structure figures were created using UCSF Chimera^58^, and the UCSF Chimera X package^64^.

### Data availability

Cryo-EM maps have been deposited in the Electron Microscopy Data Bank under accession codes: EMD-##### (CGP54626-bound GABA_B_ receptor), EMD-##### (baclofen/BHFF-bound GABA_B_ in the presence of G_i1_ protein), EMD-##### (GABA_B_-G_i_ complex in B1 state), EMD-##### (GABA_B_-G_i_ complex in B2a state) and EMD-##### (GABA_B_-G_i_ complex in B2b state). The atomic coordinates have been deposited in the Protein Data Bank under accession codes: #### (CGP54626-bound GABA_B_ receptor) and #### (baclofen/BHFF-bound GABA_B_ in the presence of G_i1_ protein).

